# Using epidemiological principles to explain fungicide resistance management strategies: why do mixtures outperform alternations?

**DOI:** 10.1101/168831

**Authors:** James A.D. Elderfield, Francisco J. Lopez-Ruiz, Frank van den Bosch, Nik J. Cunniffe

**Author notes:** Corresponding author*. Nik Cunniffe Twitter: @nikcunniffe.

## Abstract

- Whether fungicide resistance management is optimised by spraying chemicals with different modes of action as a mixture (i.e. simultaneously) or in alternation (i.e. sequentially) has been studied by experimenters and modellers for decades, largely inconclusively.
- We use previously-parameterised and validated mathematical models of wheat septoria leaf blotch and grapevine powdery mildew to test which strategy provides better resistance management, using the total yield before fungicide-resistance causes disease control to become economically-ineffective (“lifetime yield”) to measure effectiveness.
- Lifetime yield is optimised by spraying as much low-risk fungicide as is permitted, combined with slightly more high-risk fungicide than needed for acceptable initial disease control, applying these fungicides as a mixture. This is invariant to model parameterisation and structure, as well as the pathosystem in question. However if comparison focuses on other metrics, for example lifetime yield at full label dose, either mixtures or alternation can be optimal.
- Our work shows how epidemiological principles can explain the evolution of fungicide resistance, and highlights a theoretical framework to address the question of whether mixtures or alternation provide better resistance management. Our work also demonstrates that precisely how spray strategies are compared must be given extremely careful consideration.

## INTRODUCTION

Designing long-lasting, effective strategies to control plant disease remains a key challenge (Cunniffe *et al.*, 2015). Fungicide resistance management – optimising deployment to delay emergence or spread of resistant pathogen strains – has been studied for decades (Russell, 2005). Many strategies have been proposed. For a single fungicide, resistance management can be based on the method of application, changing the dose (van den Bosch *et al.*, 2011), the timing (van den Berg *et al.*, 2013), whether treatment is applied to the leaves or on the seed (Kitchen *et al*., 2016), the spatial pattern of spraying (Parnell *et al.*, 2006), or the number of sprays (van den Berg *et al.*, 2016). However – for disease control as well as resistance management – fungicides with different modes of action are often combined in a spray programme (van den Bosch *et al.*, 2014b).

Significant attention has therefore been devoted to how best to combine fungicides. Possibilities include a mixture, spraying the two fungicides at the same time; or as an alternation, applying sequentially. The risk of resistance development varies between fungicides (Brent & Hollomon, 2007). Resistance emerges to some chemicals within a few years of use, whilst others provide durable control for decades. We distinguish high-risk fungicides, to which resistance is already present or very likely to emerge, and low-risk fungicides, to which no significant resistance has yet been observed. We focus here on the case of mixture and alternation of a single high-risk fungicide with a single low-risk. Despite many experimental (Dovas *et al.*, 1976; Sanders *et al.*, 1985; Vali & Moorman, 1992; Lamondia, 2001; Cooke *et al.*, 2004) and modelling (Kable & Jeffery, 1980; Skylakakis, 1981; Josepovits & Dobrovolszky, 1985; Josepovits, 1989; Shaw, 1989a; Doster *et al.*, 1990; Birch & Shaw, 1997; Hobbelen *et al.*, 2011a, 2013) studies focusing on precisely this situation, no conclusive answer has emerged to the important but very simple question: does mixture or alternation provide better resistance management?

Although previous studies have led to equivocal results, mixtures have often been found to provide superior resistance management (van den Bosch *et al.*, 2014b). van den Bosch *et al.* (2014a) introduced a simple set of governing principles as a theoretical framework to synthesise these results, formalising previous concepts from the literature (Staub & Sozzi, 1983; Milgroom & Fry, 1988). These governing principles are based on constant rates of selection for resistance. We generalise this here, quantifying total selection for resistance by integrating a time-varying selection coefficient over time. The selection coefficient is defined as the difference in fitness between fungicide-sensitive and fungicide-resistant strains

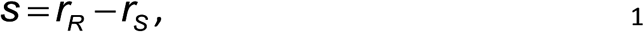

where *r*_R_ and *r*_S_ are the per capita growth rates of the resistant and sensitive pathogen strains, respectively. The total amount of selection for resistance is then given by the cumulative selection coefficient

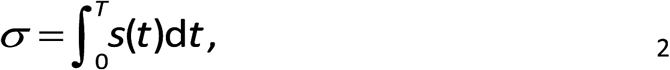

in which *T* is the time of exposure to fungicide. Selection for resistance can therefore be reduced by decreasing both *r*_R_ and *r*_S_, by decreasing *r*_R_ only, or by decreasing *T* (van den Bosch *et al.*, 2014a).

The governing principles can be applied to the comparison between mixtures and alternation. Mixtures can reduce selection as the low-risk mixing partner suppresses growth rates of both sensitive and resistant pathogens strains. Mixtures may also permit the use of less high-risk fungicide, and decreasing differences in growth rate between strains. However, due to the concave shape of fungicide dose-response curves, mixtures experience a selective cost from “dose-splitting”. Splitting a dose of fungicide over multiple sprays increases the total effect on the pathogen and thus the selection pressure imposed (van den Bosch *et al.*, 2014a). Alternations can reduce selection by reducing the number of sprays of high-risk and thus the time of exposure. Our work here assesses in detail – for the first time – the impact of this trade-off between suppression from the mixing partner and dose-splitting.

The structure of a mathematical model affects the conclusions to which it leads (Cunniffe *et al.*, 2012). Older models tended to collapse epidemics into exponential growth of fungicide-sensitive and fungicide-resistant strains (Kable & Jeffery, 1980; Skylakakis, 1981; Shaw, 1989b). Complexity has subsequently gradually increased, reflecting general trends in plant disease epidemiology (Madden, 2006), using compartmental models to represent different classes of host tissue (Gubbins & Gilligan, 1999; Hall *et al.*, 2004; Parnell *et al.*, 2005, 2006, Mikaberidze *et al.*, 2014, 2017). The current vogue emphasises detailed, system-specific models with seasonality in planting and harvesting, and a complex representation of the production of host tissue (Kitchen *et al.*, 2016; van den Berg *et al.*, 2016). What remains unclear is the extent to which divergent conclusions from previous modelling studies could be based on model structure. We therefore test how model structure affects our results. We also test the effect of pathosystem by repeating a selection of our analyses using models of two different systems. The bulk of our results are based on a model of septoria leaf blotch (caused by *Zymoseptoria tritici*) on winter wheat, but we test robustness using a model of powdery mildew (caused by *Erysiphe necator*) on grapevine. Both models have previously been parameterised and validated against field data (Burie *et al.*, 2011; Hobbelen *et al.*, 2011b).

We use these models to compare alternation and mixtures of low- and high-risk fungicides. We address the following questions.

1. Is it better to apply two fungicides as a mixture, or as an alternation?
2. Does this depend on fungicide dose, and the level of disease control?
3. How can an optimal dose and spray programme be determined?
4. Are results conditioned on: i) values of parameters governing epidemiological rates and fungicide performance; ii) model structure; and iii) the pathosystem under consideration?

The governing principles are used throughout as a unifying theoretical framework to understand the results.

## DESCRIPTION

### Modelling fungicide

#### Application strategies

We compare three strategies.

1. Mixture. Both the high- and low-risk fungicide are applied at each spray.
2. Alternation High-Low. Alternate sprays of high- and low-risk, with high-risk sprayed first in each season.
3. Alternation Low-High. High- and low-risk alternate, with low-risk first.

Since each fungicide is sprayed twice as often when part of a mixture, we halve the dose to conserve the total amount of each chemical applied per season (van den Bosch *et al.*, 2014a).

#### Fungicide dynamics

Concentrations of both fungicides are set to zero at the start of each season. The concentration of a fungicide is sharply increased whenever it is sprayed; the timing depends on the pathosystem. Between sprays there is exponential decay

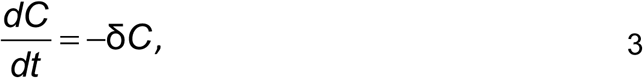

in which the decay rate δ depends on the fungicide.

#### Epidemiological effects of fungicide and dose-response

The effect of a fungicide on a pathogen depends upon its mode of action, which differs between chemicals. Protectant fungicides are assumed to affect the pathogen’s rate of infection, whereas eradicant fungicides affect the rate at which latently infected tissue becomes infectious (Hobbelen *et al.*, 2011b). Fungicides which act as a combined protectant and eradicant affect both rates.

The size of the effect upon the relevant rate parameter (ε) depends upon the fungicide’s concentration (*C*) via an exponential dose-response curve (Hobbelen *et al.*, 2011b)

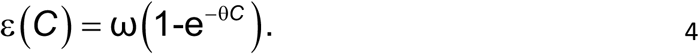

The parameters ω (maximum effect) and θ (curvature) vary between fungicides. For mixtures we assume independent action

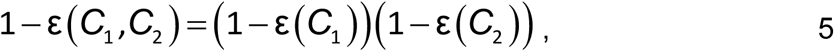

in which *C*_1_ and *C*_2_ are the individual concentrations (Hobbelen *et al.*, 2011a).

### Modelling septoria leaf blotch on winter wheat

#### Description of the model

We adapt a previously-validated, semi-discrete, compartmental model of septoria (*Z. tritici)* on winter wheat over successive growing seasons (Figure 1). Parameterisation of the model is described in Hobbelen *et al.* (2011a,b, 2013), in which the procedure for testing the fitted model against field data is also reported. The set of equations defining the model are given in Methods S1, and we concentrate here on summarising important features.

**Figure 1.**
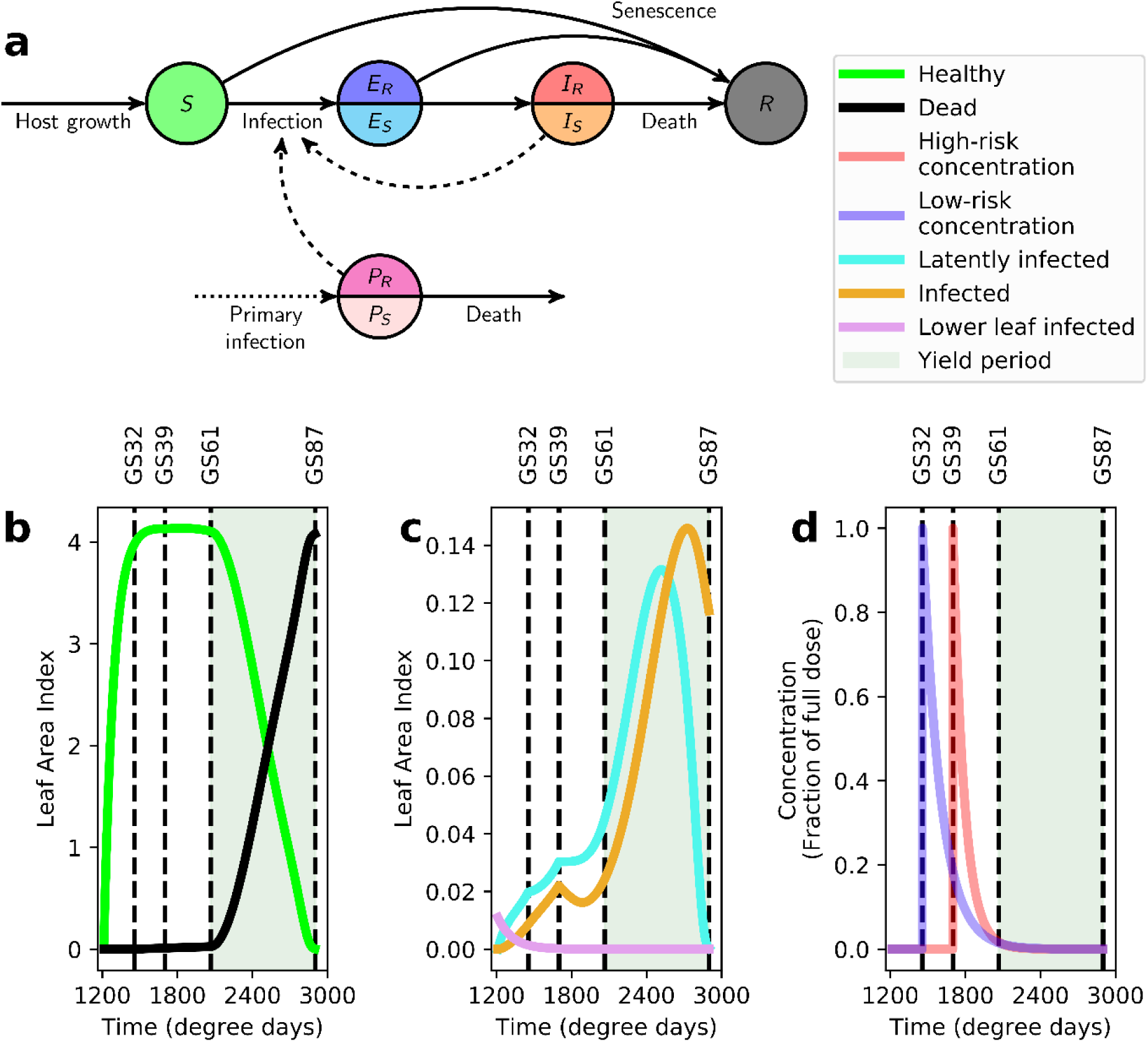
Model of septoria on winter wheat. (a) A schematic showing the structure of the within-season model for septoria on winter wheat (see also Methods S1 for the differential equations specifying the model). The model distinguishes healthy, uninfected leaf tissue (Susceptible) from different classes of infectious tissue: latently-infected (Exposed), sporulating (Infectious) and dead (Removed), as well as tracking the density of inoculum on lower leaves (Primary). Circles represent these epidemiological compartments (split into two where necessary to account for fungicide-sensitive and fungicide-resistant pathogen strains), solid lines represent transitions between compartments, dashed lines represent effects on the rates of transitions and the dotted arrow represents the point of initial infection in each growing season. Panels (b)-(d) show the dynamics of the model in the first growing season using the default model parametrisation (Table S1) and the alternation low-high strategy at full doses of both chemicals. This corresponds to a full dose of the low-risk fungicide at GS32 (1456 degree days after planting), and a full dose of the high-risk fungicide at GS39 (1700 degree days after planting). The critical time for the accumulation of yield (GS61 to GS87; 2066 to 2900 degree days after planting) is shaded; control is considered to have broken down if yields <95% of the disease-free yield are obtained. (b) The leaf area index of healthy and dead tissue over time. (c) The amount of primary inoculum and leaf area index of infected tissue over time. (d) The concentration of both fungicides over time. Note that in panels (b)-(d) dynamics start 1212 degree days after the start of the growing season; this corresponds to the time of emergence of leaf 5, with all pathogen dynamics before that time subsumed in the initial condition for the primary inoculum.

The model tracks the leaf area index (LAI), the area of leaf per unit area of ground, for the upper five leaves of wheat plants (van den Berg *et al.*, 2016). The model distinguishes healthy, uninfected leaf tissue (**<u>S</u>**usceptible) from different classes of infectious tissue: latently-infected (**<u>E</u>**xposed), sporulating (**<u>I</u>**nfectious) and dead (**<u>R</u>**emoved). The dynamics of resistance is tracked by separating fungicide-sensitive and fungicide-resistant leaf tissue (for example, splitting the infectious compartment into *I*_R_ and *I*_S_).

There are large variations in the LAI presented by a wheat crop over a single season, and this affects epidemiological dynamics (Cunniffe *et al.*, 2015). The model accounts for this by including time-dependent rates of production of healthy tissue and of natural leaf senescence. The model also accounts for decaying inoculum on lower leaves (**<u>P</u>**rimary), which initiates seasonal epidemics on the upper leaves. The amount of tissue in each compartment is reset to its initial value at the beginning of each season, with the ratio of fungicide-resistant to fungicide-sensitive inoculum set according to the corresponding ratio in the infectious compartments at the end of the previous year (or to the assumed initial frequency of resistance in the first year).

#### Fungicide effects and timing

We follow Hobbelen *et al.* (2011a) in taking pyraclostrobin as the high-risk fungicide and chlorothalonil as the low-risk fungicide. We assume that pyraclostrobin acts as a combined protectant and eradicant, but that chlorothalonil has only protectant activity. We consider two applications of fungicide per season, with a T1 spray at Zadoks growth stage 32 and a T2 spray at growth stage 39. This is representative of those used in winter wheat growing areas (Paveley *et al.*, 2014).

#### Calculating yield

We estimate the relative yield (*Y*) by integrating the amount of photosynthetically-active leaf area over a critical period for grain formation (Waggoner & Berger, 1987; Gooding & Dimmock, 2000)

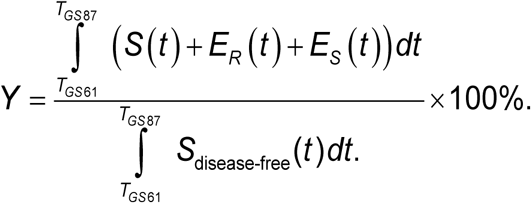

The denominator normalises the yield to that obtained from a disease-free crop.

### Strategy performance

The goal of any anti-resistance strategy is maintaining effective disease control. However, this begs a question: what level of control is effective? We define a threshold level of disease beyond which management is considered to have failed, taking a 5% yield loss as the critical level growers will tolerate (Hobbelen *et al.*, 2011a). The effective lifetime of the high-risk fungicide (the “usefulness time” of van den Bosch & Gilligan, (2008)) is defined as the number of seasons until this critical yield loss occurs.

We use the following metrics.

- **Selection ratio (SR)**. Proportional increase in the frequency of resistance over the first season, i.e. the proportion of the total infectious tissue (*I_R_* + *I_S_*) infected by the resistant strain (*I_R_*). This measures the rate at which fungicide-resistance spreads initially.
- **Lifetime yield (LY)**. Total within-season yield over the entire effective lifetime. This allows strategies that have similar effective lives but differences in within-season performance to be distinguished.

Comparison of strategies depends on the metric. For selection, we define *Z* = *SR*_ALT_/ (*SR*_ALT_ + *SR*_MIX_), in which *SR*_ALT_ is the selection ratio of the best-performing of the two alternation strategies, and *SR*_MIX_ is that of mixtures. Since smaller values of the selection ratio are superior, values of *Z* lower than 0.5 indicate alternation is preferred. For lifetime yield, larger values indicate a better performing strategy, and we preserve the directionality of the *Z*-metric by instead defining *Z* = *LY*_MIX_ / (*LY*_ALT_ + *LY*_MIX_), again using the best-performing alternation strategy in the comparison.

### Effect of model structure

We check the robustness of our results to the set of mechanisms included in the underlying epidemic model. We identify three components that could be significant.

1. **Host-limited infection**. The density of host tissue is modelled in some detail, with a complex time-dependent function representing production of susceptible host tissue, whereas simpler models often use exponential growth.
2. **Latent period**. New infections only become infectious after a latent period, whereas in simpler models infected tissue becomes infectious immediately.
3. **Phenology**. The model includes a complex treatment of within-season timing, with primary inoculum from lower leaves initiating upper leaf epidemics, and also the senescence of living leaf tissue. These features are absent from the simpler models.

These complexities are sequentially taken out of the model and analyses re-run (Methods S2).

### Effect of pathosystem

#### Modelling powdery mildew on grape

As a further test of robustness, we repeat a selection of analyses using a model of powdery mildew (*E. necator*) on grapevine (Burie *et al.*, 2011) (Figure 2). In addition to being parameterised to match the grapevine powdery mildew pathosystem, there are three additional differences in model structure in comparison to the septoria model.

**Figure 2.**
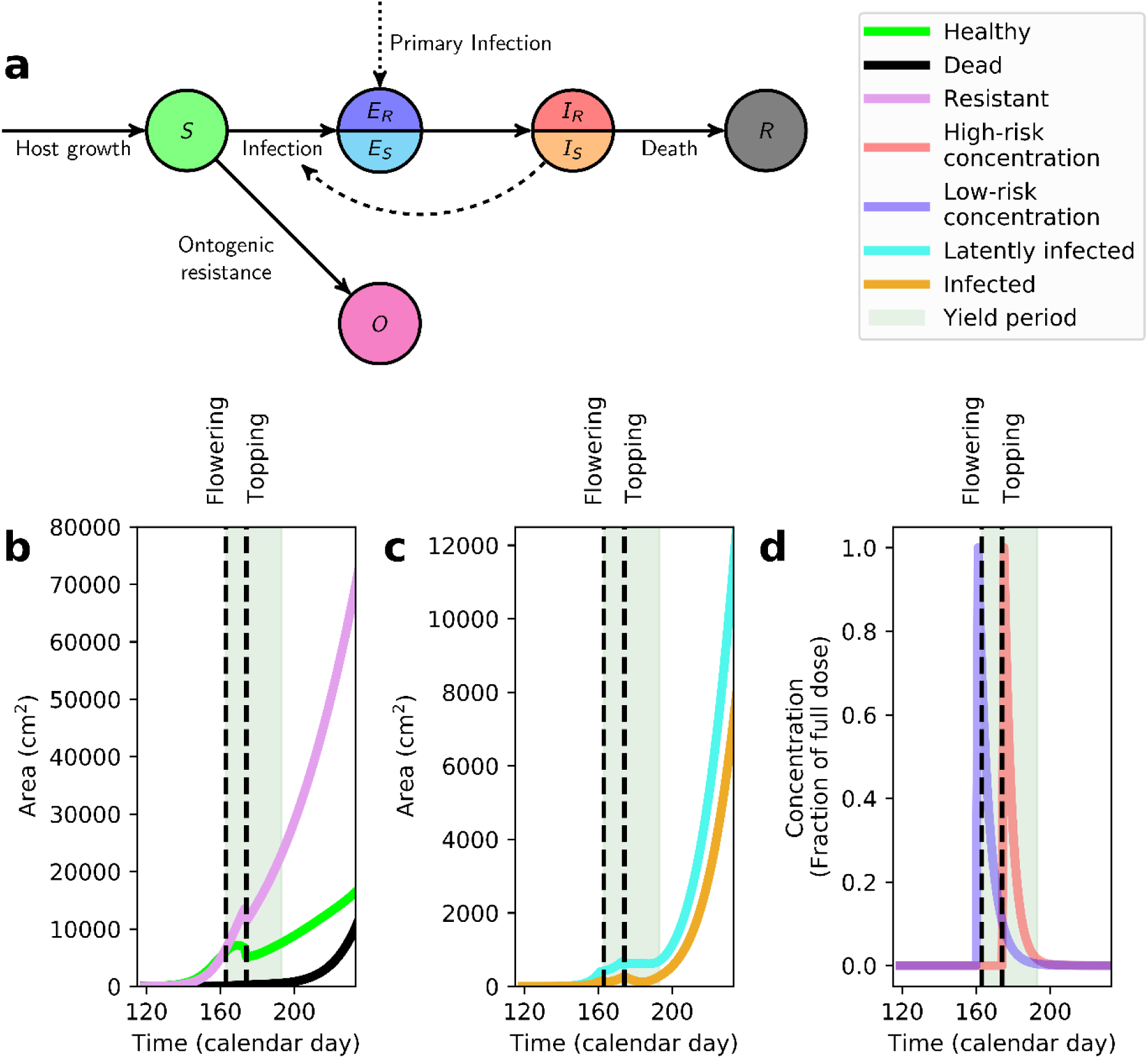
Model of powdery mildew on grapevine. (a) A schematic showing the structure of the within-season model for powdery mildew on grapevine (see also Methods S3 for the differential equations specifying the model). The model distinguishes healthy, uninfected leaf tissue (Susceptible) from different classes of infectious tissue: latently-infected (Exposed), sporulating (Infectious) and dead (Removed), as well as (Ontogenic) tissue which has become resistant to infection by virtue of its age. Again, circles represent epidemiological compartments (split into two where necessary for each pathogen strain), solid lines represent transitions between compartments, dashed lines represent effects on the rates of transitions and the dotted arrow represents the point of initial infection in each growing season. Panels (b)-(d) show the dynamics of the model when a full dose of the low-risk is applied at the first spray (at day 161,2 days before flowering occurs at day 163) and of the high-risk at the second spray (day 175, 12 days after flowering). Control is considered to have broken down when the severity of disease exceeds 3% within the period from flowering to 30 days later. (b) The area of healthy and dead tissue over time. (c) Area of infected tissue over time. (d) Concentration of fungicide over time. Note the agronomic practice of topping is modelled as being performed 10 days after flowering (day 173), and this leads to sharp changes in the values of state variables and epidemiological parameters at this time (Methods S3).

1. There is no primary inoculum compartment, and epidemics are instead initiated by a small amount of tissue being set to be latently infected (i.e. exposed) at the start of each season.
2. An additional compartment is included in the model, accounting for leaves developing **<u>O</u>**ntogenic resistance by virtue of age.
3. The model includes shoot-topping, in which upper shoots are removed to encourage secondary shoot growth.

Full details of the model – and its parameterisation – are in Methods S3.

#### Fungicide effects and timing

For powdery mildew we model trifloxystrobin as the high-risk fungicide, and sulphur as the low-risk fungicide, assuming both chemicals combine protectant and eradicant modes of action (Reuveni, 2001). We assume flowering occurs at day 163 of the season (Mammeri *et al.*, 2014), and that spraying is done either side of this, two days before and twelve days after flowering. This is a smaller number of sprays than normally used in French viticulture (Calonnec *et al.*, 2006; Savary *et al.*, 2009), although it is within the range leading to acceptable control (Gadoury *et al.*, 2003).

#### Effective lifetime and yield

The Burie *et al.* (2011) model tracks the severity of powdery mildew on grapevine leaves. However, prices obtained by a grower would depend on a combination of yield and grape quality for winemaking. Effects of leaf infection upon yield and quality are complex (Pool *et al.*, 1984; Calonnec *et al.*, 2004). Quantifying fine details of this would require a more detailed treatment than appropriate here. However, there is a strong positive correlation between leaf infection and berry infection (Calonnec *et al.*, 2006; Delière *et al.*, 2015). We therefore simply use the level of leaf infection as a proxy for yield, taking 3% as the critical threshold on the peak level of berry infection within 30 days of flowering beyond which control is considered to have broken down, since berries are almost entirely resistant after this period (Gadoury *et al.*, 2003). Similarly stringent thresholds are used in French viticulture (Deliere *et al.*, 2010). We then set the equivalent of lifetime yield to be the effective lifetime of the high-risk fungicide, i.e. the number of seasons until this critical threshold is exceeded.

## RESULTS

### Initial disease control at full doses

For septoria and applying full doses of both fungicides, all three strategies lead to adequate control in the first season (yield >95% of the disease-free yield) (Figure 3a). Optimal initial control is obtained under mixture (~97.4% yield), with lower yields from both alternation strategies (~96.0% and ~96.4% for low-high and high-low, respectively). The first-season yield is highest for mixtures because of the concave dose response curve, with diminished returns from increased concentrations. Spraying half the dose twice as often therefore leads to better control (recall full dose corresponds to half dose in both sprays under mixture). The alternation high-low strategy slightly outperforms the low-high strategy in the first season since the high-risk fungicide is assumed more efficacious (maximum effect ω_H_ = 1.0 > 0.48 = ω_L_). All other things being equal, control is improved by applying the high-risk fungicide earlier, since it then targets the pathogen when its relative growth rate is larger.

**Figure 3.**
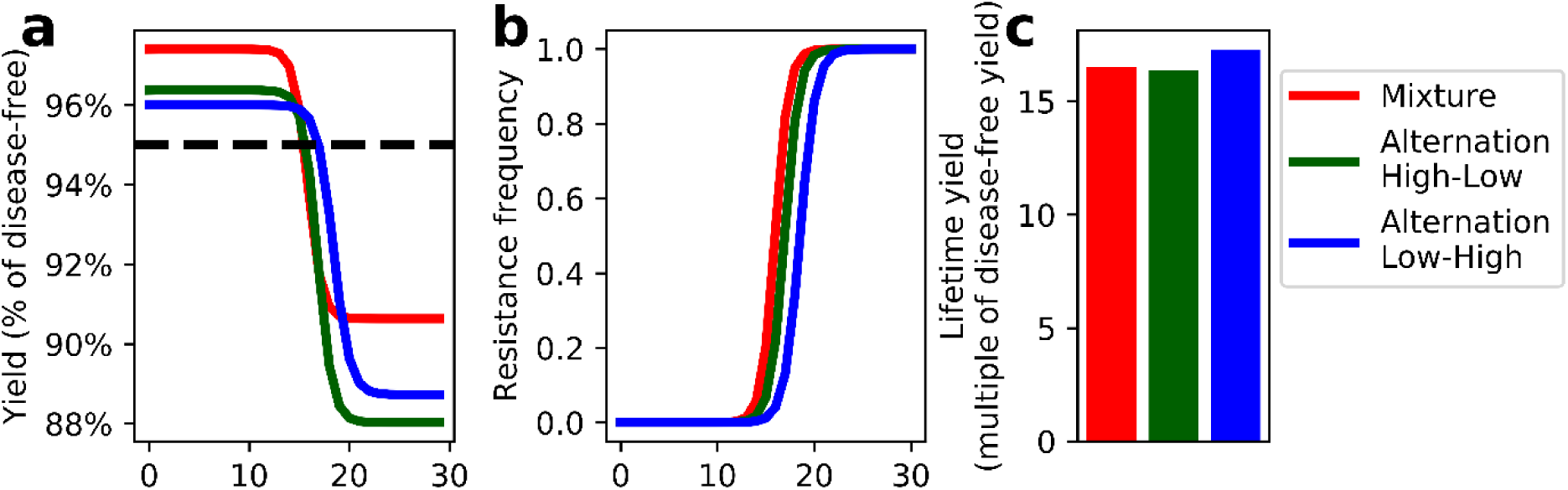
Performance at full doses of both fungicides. (a) The yield – expressed as a percentage of the disease-free yield – as a function of the growing season, for all three strategies at full doses. The dashed line corresponds to the critical yield threshold (95% of the disease-free yield) below which control is assumed to be economically ineffective. (b) The frequency of resistance to the high-risk fungicide in the pathogen population at the start of each growing season. (c) The overall lifetime yield (expressed as a multiple of the disease-free yield in a single season).

### Evolution of resistance at full doses

For all three strategies, there is a sharp breakdown of control after ~15 seasons (Figure 3a). This is driven by a rapid increase in the proportion of the resistant strain, which increases sigmoidally from being practically undetectable (<1%) to near fixation (>99%) within one or two growing seasons (Kable & Jeffery, 1980) (Figure 3b). Disease control then rests entirely on the low-risk fungicide, and all three strategies become ineffective (i.e. yield <95%). Due to dose-splitting and the concave dose-response curve, when resistance is at high frequency the best yield is then obtained under mixtures (Figure 3a), although this level of control is not economically-viable. The improved performance of the low-high strategy relative to high-low alternation after resistance has taken over is again due to timing: control is improved by applying the sole effective fungicide earlier.

Although the timing of the sharp increase in the frequency of fungicide-resistant pathogen is similar for all three strategies, it occurs earliest for mixtures then for alternation high-low then for low-high (Figures 3a and b). This is precisely the order of the efficacy of the strategies for disease control in the first season. Applying fungicides as a mixture leads to slightly more effective disease control, but – in part as a consequence of this – exerts a stronger selective pressure. Considered over the effective lifetime, the alternation low-high strategy therefore has the highest lifetime yield (Figure 3c). At full dose, however, differences between the strategies are relatively minor.

### Responses of selection and lifetime yield to dose

Full dose results illustrate disease control and selection are closely related. We therefore consider performance over a range of dose combinations, identifying how to select a pair of doses – as well as spray strategy – to optimally balance control *vs.* selection. For selection, there are dose combinations favouring both alternation and mixtures (Figure 4a). Alternation exerts less selection than mixtures at higher doses of high-risk (*C_H_)*. The concave dose-response means that at high doses the effect on pathogen growth rates of the half dose under mixture approaches that of full dose under alternation. However, under mixture this dose is applied twice as often, and selection occurs for longer. Conversely, at higher doses of the low-risk fungicide (*C_L_)*, mixtures tend to be preferred, since alternation receives no suppression of pathogen growth rate from the low-risk at the time the high-risk fungicide is applied.

**Figure 4.**
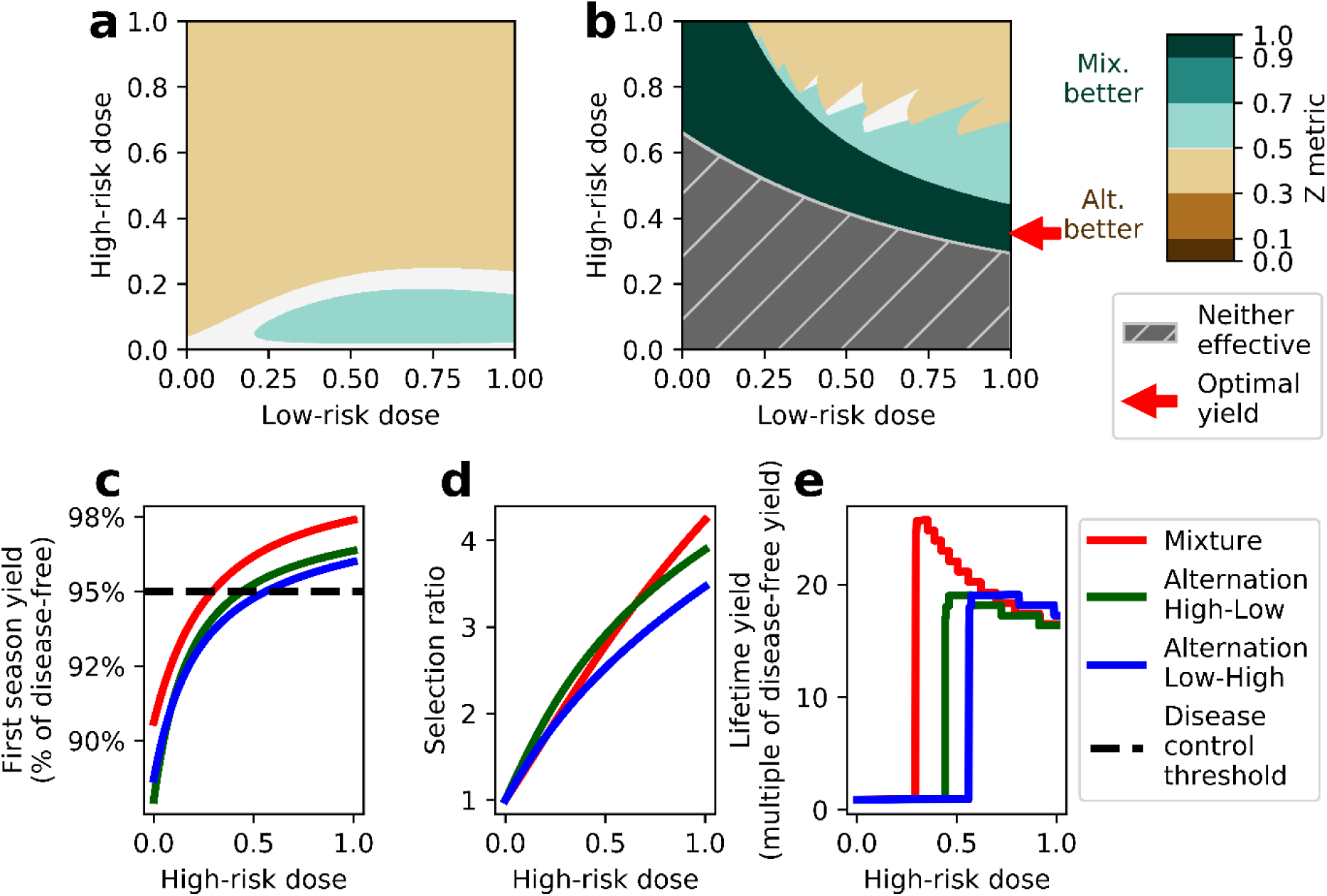
Defining the optimal spray strategy and dose combination. (a) Comparison of the selection ratio (*SR*) for mixtures *vs.* alternation, for different doses of high-risk and low-risk fungicides (over the entire growing season, and so under mixtures each dose as shown in dose-space is halved at the time of application; this is done for all results presented in this paper). In all cases the selection ratio under mixtures is compared with the best-performing of the two alternation strategies. The Z metric – which is used to visualise the comparison – is defined here as *SR*_ALT_ / (*SR*_ALT_ + *SR*_MIX_). Values lower than 0.5 therefore indicate alternation is superior, and values greater than 0.5 indicate mixtures are superior. Values very close to 0.5 – which are shaded white – correspond to the two strategies performing equally well. (b) Comparison of the lifetime yield (*LY*) for mixtures *vs.* the best performing alternation strategy. The Z metric is defined here as *LY*_MIX_ / (*LY*_ALT_ + *LY*_MIX_): again higher values indicate mixtures are superior. Regions of dose-space within which no spray strategy can provide effective control in the first growing season are hatch-shaded grey; the dark green region corresponds to areas of dose-space within which disease can only be controlled when the fungicides are sprayed as a mixture. The largest lifetime yield is marked with the red arrow, and occurs when spraying using a mixture including the maximum dose of low-risk fungicide (*C_L_* = 1.0). Panels (c)-(e) show results for all three strategies as a function of high-risk dose when the low-risk dose is fixed at *C_L_* = 1.0. Individual panels: (c) first-season yield; (d) selection ratio; and (e) lifetime yield. In (c) the dashed line shows the disease control threshold.

Patterns in lifetime yield are more complex (Figure 4b). There is a region of dose-space (hatch-shaded grey) within which no strategy leads to sufficient control even before resistance has spread. This outcome is associated with low doses of both chemicals, although even at full dose of low-risk, some high-risk is required (recall the high-risk chemical is the more efficacious). There is an intermediate region (shaded dark green) within which – at the same doses of both chemicals for each strategy – effective control is only possible under mixture because of dose-splitting. For larger doses of both chemicals either mixtures (shaded light green) or one of the alternation strategies (shaded light brown) can have the best performance, or there can be approximately equal lifetime yields (within 1%; shaded white).

### Selecting an optimum strategy and dose

The largest lifetime yield over all strategies and pairs of doses is marked with the red arrow on Figure 4b. It corresponds to spraying a mixture of a full dose of low-risk with a dose of high-risk slightly larger than that required for economically-acceptable yield in the first season. As the low-risk fungicide exerts no selection, it is unsurprising that the maximal permissible amount of low-risk should be optimal, since this allows the smallest amount of high-risk to be applied while maintaining acceptable disease control. However, it is less obvious why the optimal strategy should be to apply the high-risk fungicide as part of a mixture, and why this particular dose of high-risk (i.e. just above the amount required to ensure effective control) is required.

We therefore examine responses to the dose of high-risk (*C*_H_) with the low-risk fixed at full dose (Figures 4c-e). This corresponds to the vertical line in dose-space *C_L_* = 1.0 in Figures 4a and b. As already noted, for a given value of *C*_H_, mixture leads to the best initial disease control because of the beneficial effect of dose-splitting (Figure 4c). The implication is that lower *C*_H_ can maintain adequate control under mixtures (*C*_H_ ≥ 0.29; dashed line in Figure 4c) compared to alterations (*C*_H_ ≥ 0.44 or 0.56). The pattern for the level of selection to *C*_H_ is more complex, with both mixture and alternation potentially leading to smaller selection ratios at different *C*_H_ (Figure 4d).

To understand optimum performance in more detail, we compare the selection ratios at the low end of permissible doses for each strategy, since these maximise lifetime yields (Figure 4e). For the model and parameterisation used here, the lower permissible dose under mixture outweighs the effect of spraying high-risk twice as often, and exerts less selection than the lowest permissible doses under either alternation (at 95% yield in the first season, SR = 2.28 for mixture *vs.* 3.08 and 3.15 for alternation high-low and low-high, respectively). The lower selection ratio leads to a longer effective lifetime, and spraying the fungicides as a mixture optimises lifetime yield.

The optimal dose of the high-risk fungicide (*C_H_*) is slightly higher than the minimum *C_H_* ensuring acceptable control in the first season. This is because the effective lifetime is discrete, leading to ranges of *C_H_* which all break down within the same season. Within any range of doses with the same effective lifetime, the optimum lifetime yield is obtained by selecting a higher *C_H_*, benefitting from slightly improved control in each season it remains effective. Too high a *C_H_* however can lead to more dramatic failure in the final season, and thus the optimal *C_H_* may not be the highest dose with the longest effective life. This is difficult to see in Figure 4e, but the “horizontal” parts of the response are not quite horizontal. The red arrow on Figure 4b is therefore above the boundary between the grey and dark-green regions.

### Balancing selection and control

We further dissect the trade-off between selection and control by considering equal doses of high- and low-risk (Figures 5a-e), corresponding to a different visualisation of results underpinning the line *C_H_* = *C_L_* in Figures 4a and/or 4b. For all three strategies, as the dose of high- (and low-) risk is increased, both the first season yield (Figure 5a) and selection ratio (Figure 5b) increase. However, dose-splitting means that viable control is again retained under mixtures at much lower doses. The overall maximum lifetime yield is therefore under mixtures (*LY* = 19.1, *C_L_* = *C_H_* = 0.5) (Figure 5c). Strategies can be normalised against each another by replotting the selection ratio and lifetime yield as a function of the first season yield (Figures 5d and e). This reiterates that – at least at the same level of initial disease control and using equal proportions of both chemicals – mixtures leads to less selection (Figure 5d) and a larger lifetime yield (Figure 5e). Differences between strategies are small, however.

**Figure 5.**
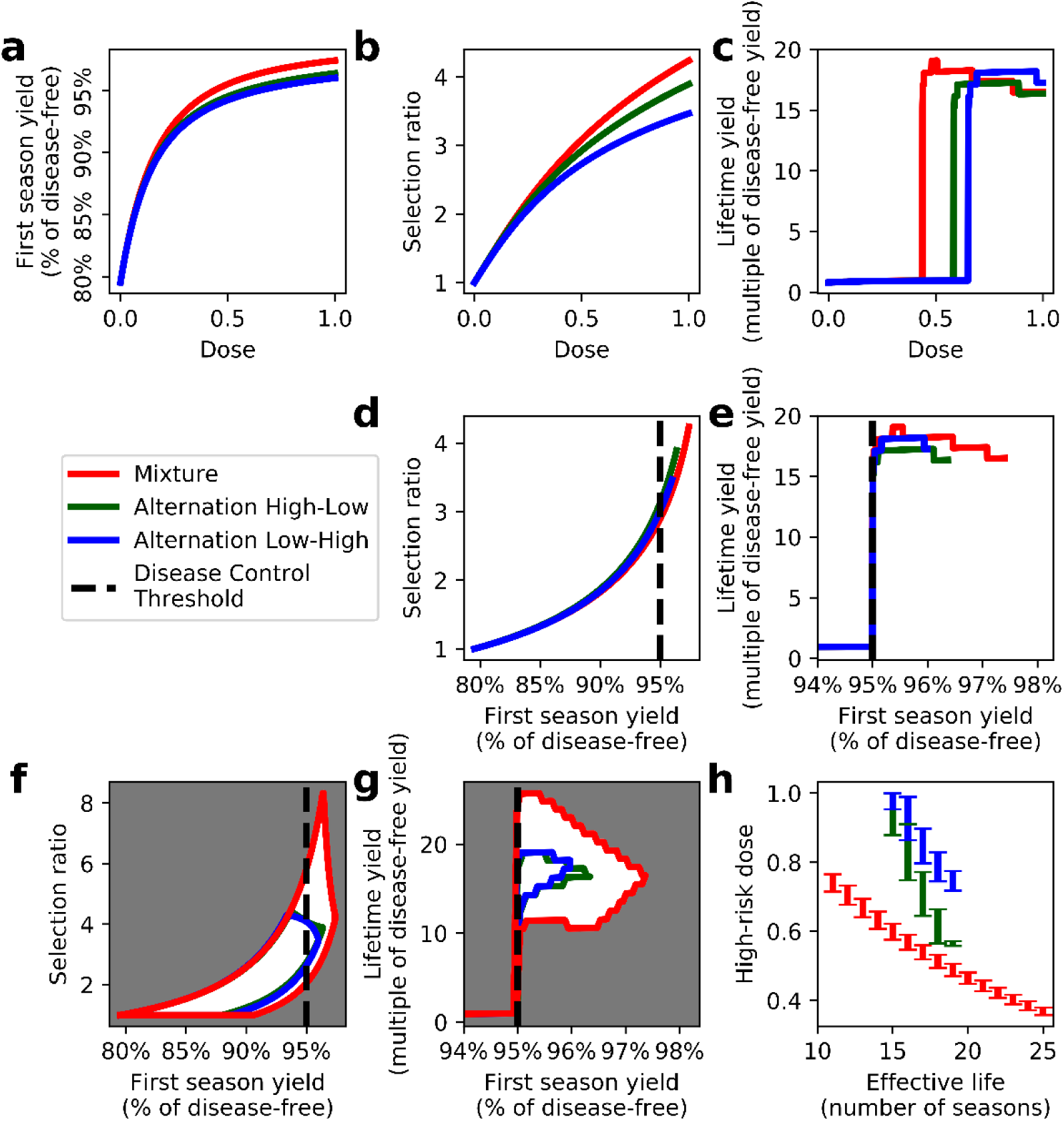
Balancing selection and disease control. Panels (a)-(c) show performance as the dose varies but when equal amounts of low- and high-risk fungicide are sprayed (i.e. *C_L_* = *C_H_*). Individual panels are: (a) first-season yield; (b) selection ratio; and (c) lifetime yield. Panels (d) (selection ratio) and (e) (lifetime yield) show the results from (b) and (c) plotted as a function of the first season yield. When normalised for the level of initial disease control, mixtures lead to less selection and larger lifetime yields than either alternation strategy. Panels (f)-(h) show the results when the constraint that doses of both fungicides should be equal is removed. For all three strategies there are ranges of values of the selection ratio and the lifetime yield that correspond to each single value of the first season yield, with different relative proportions of highrisk to low-risk fungicide that is sprayed. The top and bottom of the ranges of the selection ratio and the lifetime yield are shown for each strategy, although all intermediate points can be attained for different combinations of low- and high-risk chemicals. The ranges are wider for mixtures than for either alternation strategy, meaning that – depending on the dose of high-risk fungicide – mixtures can lead to both better and worse outcomes for resistance management at any level of disease control. However, dose combinations that cause mixtures to provide the most effective resistance management can always be selected. (h) The range of high-risk doses that lead to different effective lifetimes, for each strategy (first season yields between 95.45% and 95.55%; this corresponds to vertical slice through the data shown in panel (g)). A wider range of effective lifetimes are possible at this level of disease control under mixtures, and a given effective lifetime results from a smaller dose of high-risk fungicide. In (a), (d), (e), (f) and (g) the dashed line shows the disease control threshold.

Examining responses of selection and control to first season yield is a convenient mechanism to allow results at all doses – not just when *C_H_* = *C_L_* – to be visualised (Figures 5f-h). For any given initial level of control, mixtures can produce higher lifetime yields and lower selection ratios (Figures 5f and 5g). However, mixtures can also produce lower yields and higher selection ratios. The variation in selection and yield for a given level of control is therefore much larger with mixtures than with either alternation. Examining strategies that have the same initial disease control shows that mixtures can produce any particular effective lifetime at much lower *C_H_*. This is shown in Figure 5h, which – for the ranges of doses of both fungicides under all three strategies which lead to first season yields between 95.45% and 95.55% – shows values of *C_H_* leading to each effective lifetime (i.e. a vertical slice through the data underpinning Figure 5g).

### Effect of epidemiological and fungicide parameters

Results thus far correspond to a single model parameterisation. We test robustness by altering values of a number of key parameters. In all cases the dose of low-risk fungicide is fixed to be maximal (i.e. *C_L_* = 1.0), as we have identified no mechanism by which changing parameter values can cause this not to be optimal. We then consider the response of lifetime yield to changing *C_H_*.

As an example, we examine in some detail the effect of the infection rate (β) (Figure 6a). If β is made significantly larger than the default, all three strategies fail to give sufficient control at any *C_H_* (dark-grey hatching). If β is made sufficiently smaller, then control can be maintained indefinitely through the low-risk fungicide alone (light-grey hatching). Both cases are unrealistic. Within the realistic range of values of β, exactly the same pattern is seen as before. At low *C_H_*, no spray strategy can provide effective control. At slightly higher *C_H_* mixtures perform best. At the highest *C_H_* alternation performs better, although for large infection rates this might require *C_H_* > 1.0 (i.e. a dose above the permissible maximum label dose). As β is increased, the threshold *C_H_* at which mixtures first become effective shifts upwards, as more fungicide is required to provide acceptable disease control.

**Figure 6.**
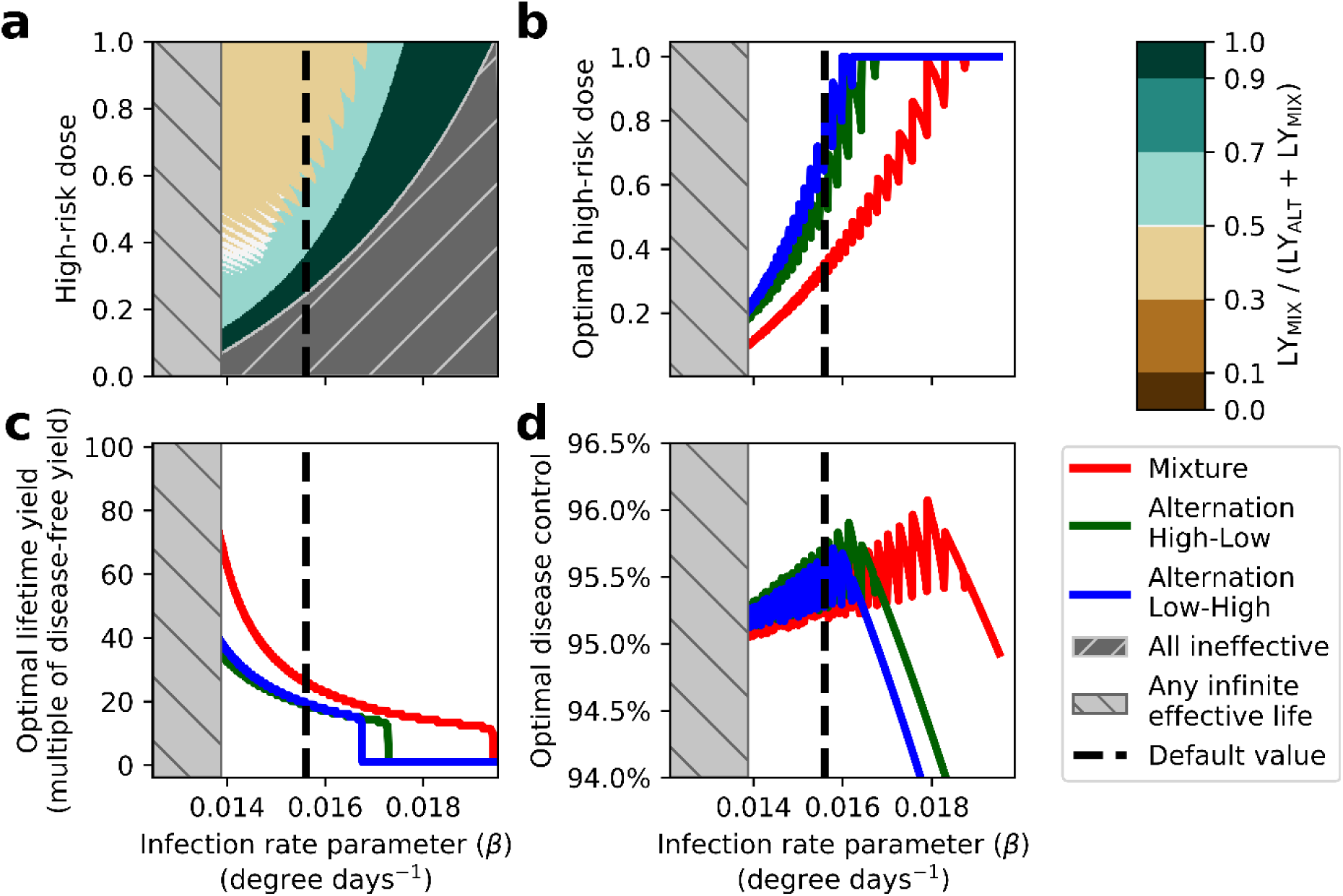
Response to the infection rate parameter. (a) The value of the Z metric for lifetime yield (i.e. Z = *LY*_MIX_ / (*LY*_ALT_ + *LY*_MIX_)) for different values of the infection rate β (x-axis) and doses of high-risk chemical, *C_H_* (y-axis), when the dose of the low-risk chemical is fixed at *C_L_* = 1.0. The dark grey region corresponds to pairs of infection rates and high-risk doses for which no strategy can provide effective control in the first growing season, and the light grey region corresponds to at least one of the strategies providing effective control without any high-risk chemical being sprayed (and so within which resistance management is trivial). (b) The optimal dose of high-risk chemical for each strategy as a function of the infection rate. (c) The corresponding lifetime yield that is obtained. (d) The level of disease control – as measured by the yield in the first season – at optimum for each strategy. In panels (b) and (d) the saw-tooth pattern is because the effective lifetime – the most important determinant of the lifetime yield – is a discrete quantity. The dashed lines in all panels shows the default value of β.

The optimal dose is always lower for mixtures than either of the alternations (Figure 6b; the saw-tooth pattern is because the effective lifetime is discrete). The corresponding optimal lifetime yield is always larger under mixture (Figure 6c), and – for all strategies – corresponds to selecting *C_H_* close to the threshold required for effective first season control (Figure 6d). For all values of the infection rate, β, the optimal strategy is therefore again to spray a little more high-risk fungicide than required for effective control in the first season, and to do so under mixture.

The pattern is consistent for all parameters tested in our sensitivity analysis (Figure 7). For parameters which cause disease to spread faster as they are increased, more high-risk fungicide is required for effective control, and the characteristic pattern shifts upward (Figures 7c and f). Conversely, for parameters for which an increase leads to decreased rates of disease spread, a smaller amount of high-risk is permissible (Figures 7a, b, d, e and g). Changing the initial frequency of resistance has a negligible effect on the relative performance of the strategies in the first season, for both resistance management and yield. However at higher initial levels of resistance, control fails sooner, increasing the importance of disease control in the earlier seasons for the lifetime yield and thus favouring mixture (Figure 7h).

**Figure 7.**
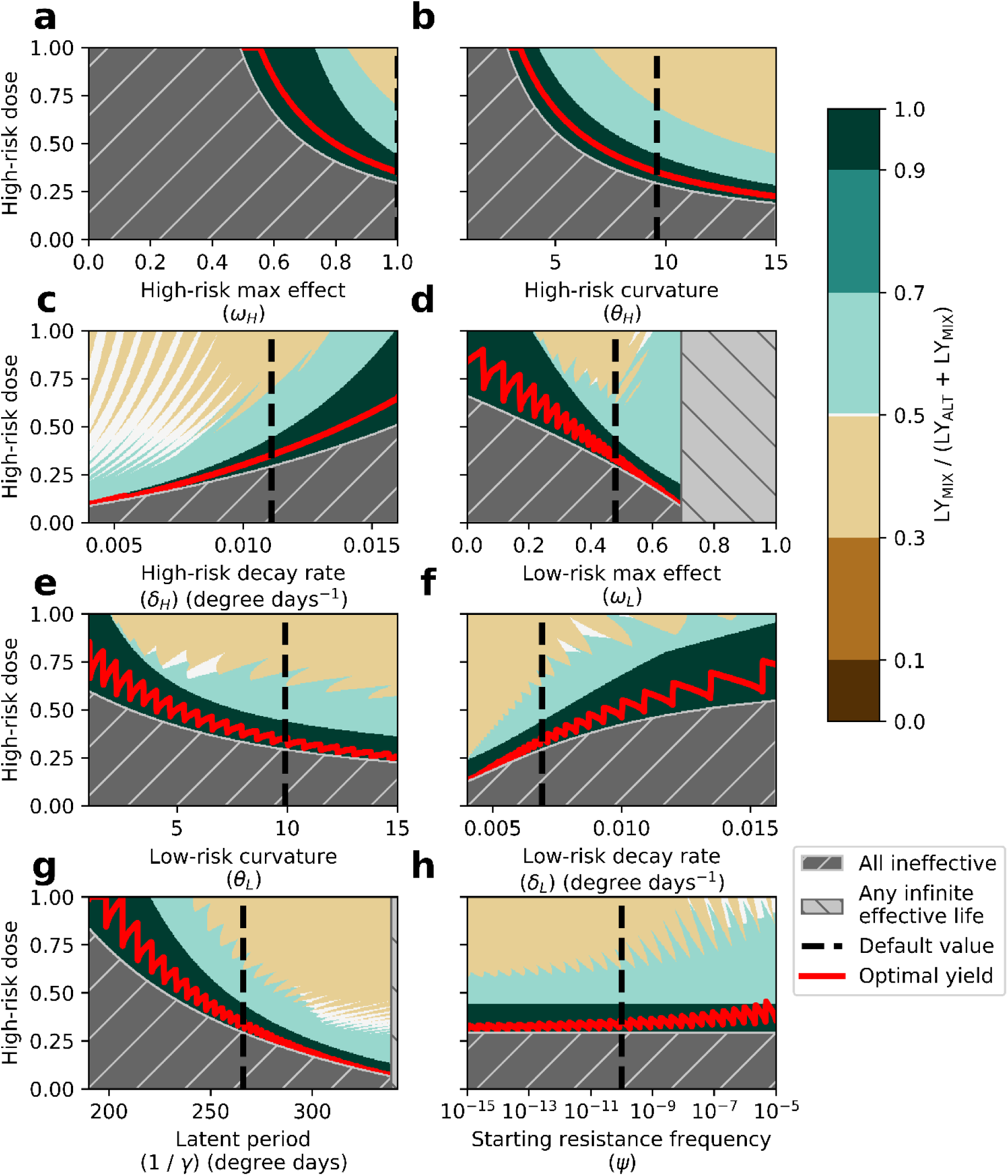
Full sensitivity analysis for a range of epidemiological and fungicide parameters. Each panel shows the equivalent of Figure 6(a) for a different parameter. (a) Maximum effect parameter for the high-risk fungicide. (b) Curvature parameter for the high-risk fungicide. (c) Decay rate for the high-risk fungicide. (d) Maximum effect parameter for the low-risk fungicide. (e) Curvature parameter for the low-risk fungicide. (f) Decay rate for the low-risk fungicide. (g) Pathogen latent period. (h) Initial frequency of the resistant pathogen strain. The red lines mark the optimal doses of high-risk fungicide for lifetime yield for each value of the parameters: in all cases this corresponds to applying the two fungicides as a mixture.

### Effect of model structure

To facilitate inter-model comparison, we return to comparing strategies in dose-space. The simplest model – with both pathogen strains growing exponentially – is similar to models used in the early fungicide resistance modelling literature (Delp, 1980; Kable & Jeffery, 1980; Skylakakis, 1981). Indeed, if we additionally assume fungicides do not decay, an analytical prediction of which strategy leads to better resistance management at a given pair of doses can be generated (Methods S4). The other models are too complicated for mathematical analysis, although the same pattern is seen for selection in dose-space in every model (Figure 8). The only real differences between models are the slightly larger regions within which alternation provide better resistance management when models include a latent period. When a latent period is not included, the high-risk fungicide loses its eradicant mode of action, and so becomes generally less efficacious. As with dose and fungicide parameter values, less effect from the high-risk fungicide then favours mixture.

**Figure 8.**
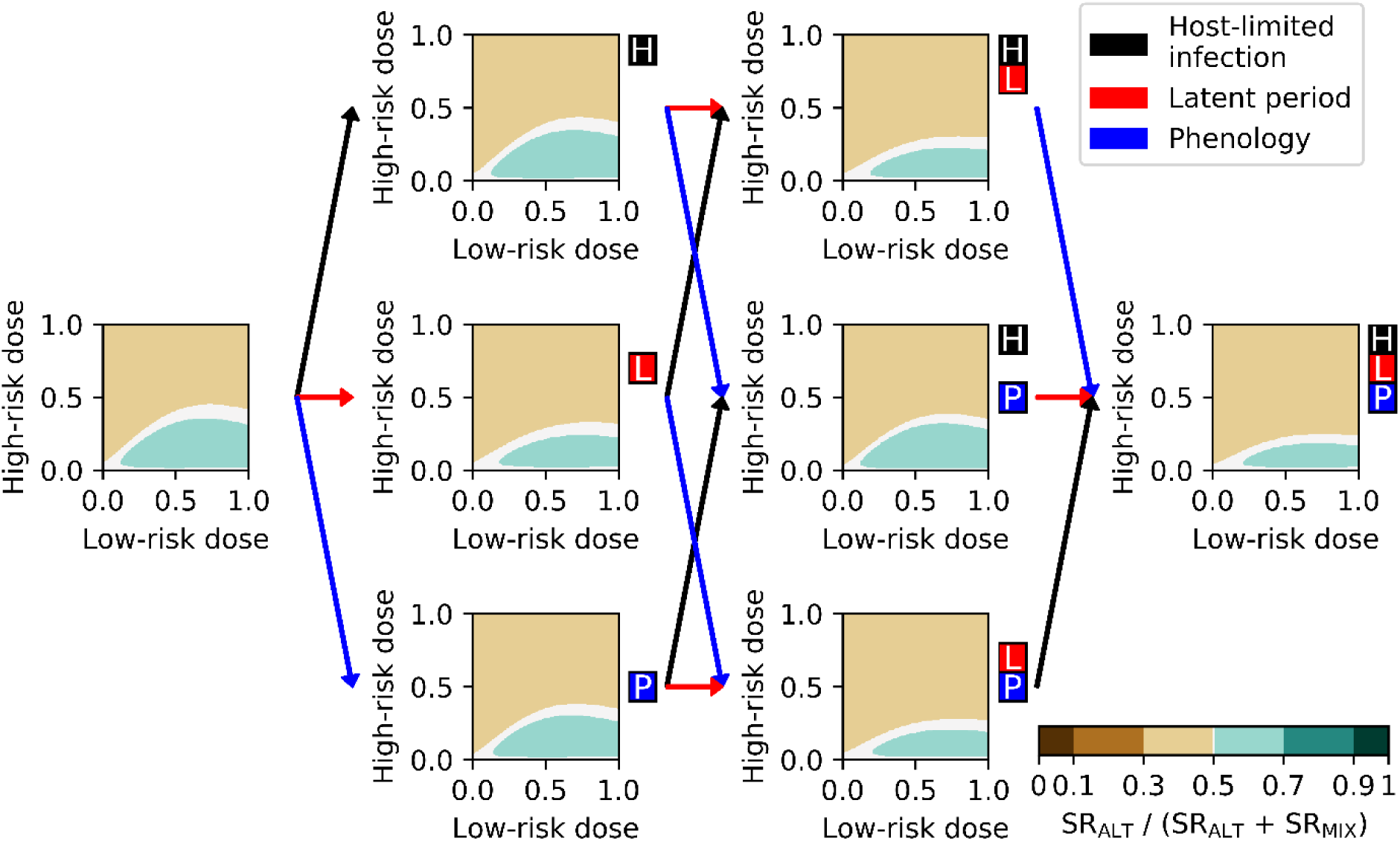
Effect of model structure on the comparison between mixtures and alternation for selection. The relative performance of mixtures and (the best-performing) alternation is plotted in dose-space, by calculating the Z metric, Z = *SR*_ALT_ / (*SR*_ALT_ + *SR*_MIX_), for all of the powdery mildew sub-models investigated. The coloured boxes next to each subplot identify what model features are present, and the coloured lines show which features differ between connected sub-models. The right-most model includes all features, and so corresponds to the full model of septoria considered in the bulk of the paper (i.e. the right-most plot is exactly as Figure 4a).

In the models that include host-limited infection, and thus the loss of host tissue to disease, we also investigate how predictions of lifetime yield are affected by model structure (Figure 9). Predictions vary between models, which is perhaps unsurprising given the additional complexity underlying the yield metric. However, although the patterns vary, the characteristic pattern in dose-space is conserved. At low *C_H_* both strategies fail to give acceptable yield, at slightly higher doses mixtures out-perform alternation, and at the highest doses alternation out-performs mixture. Exactly as before, the optimal strategy is therefore again to spray a little more high-risk fungicide than is required for effective control in the first season, and to do so under mixture (the bright green saw-tooth lines on Figure 9).

**Figure 9.**
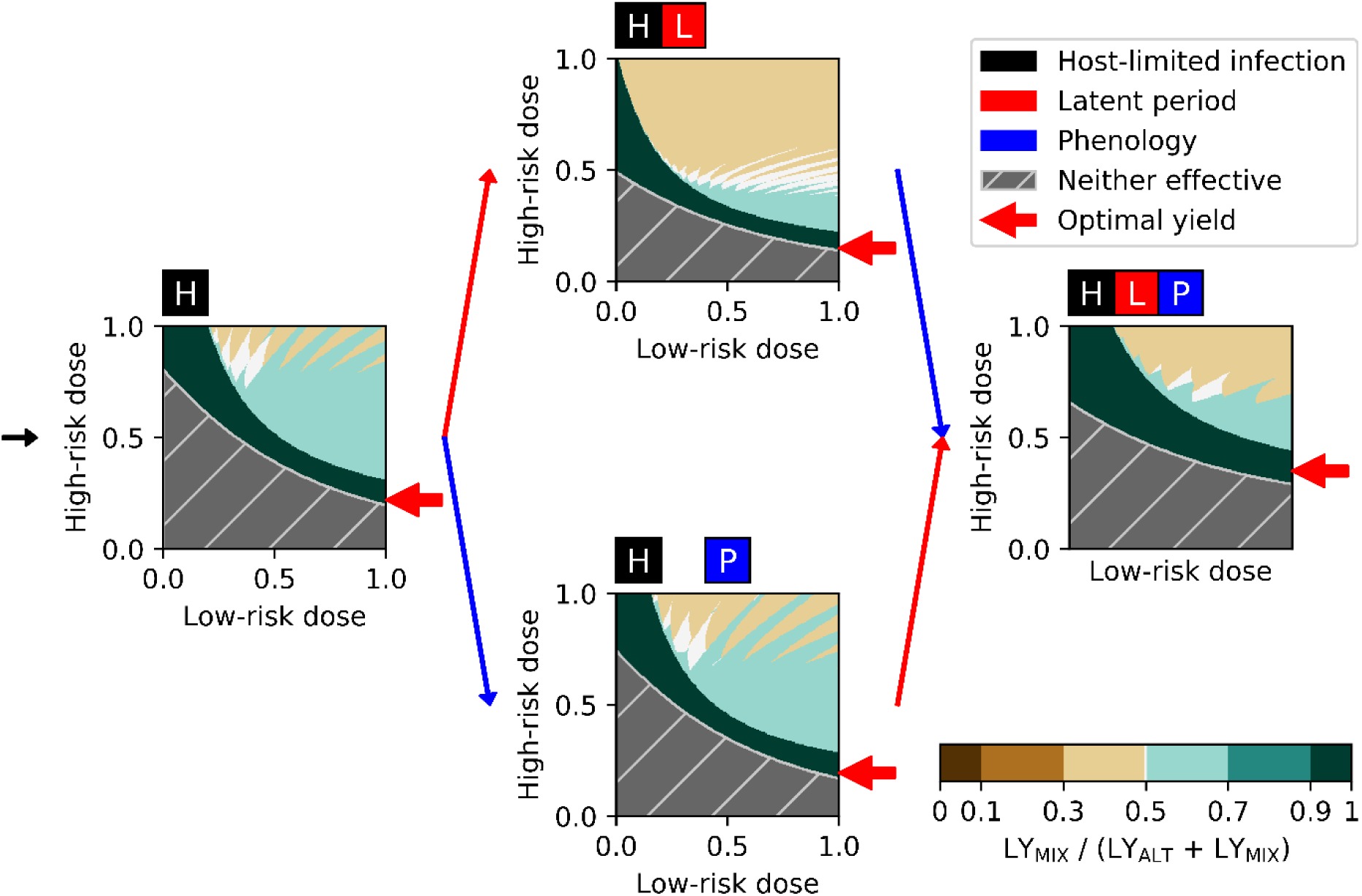
Effect of model structure on the comparison between mixtures and alternation for lifetime yield. The relative performance for lifetime yield of mixture and (the best-performing) alternation is plotted in dose-space, showing Z = *LY*_MIX_ / (*LY*_ALT_ + *LY*_MIX_), for a range of the powdery mildew sub-models investigated. Note that it does not make sense to consider the yield in models which do not contain host limitation, and so these models are omitted. The optimal lifetime yield is marked by the red arrow (in all cases this is at full-dose of low-risk and is obtained under mixture). Note again that the right-most model corresponds to the full model, and so replicates Figure 4b.

### Effect of pathosystem

Results for powdery mildew are similar to those for septoria for both comparisons in dose-space, with alternation performing better at higher doses of the high-risk fungicide in terms of both resistance management and long-term yield (Figures 10a and 10b). Mixtures perform increasingly well for selection if *C_L_* is increased, while the yield metric produces more complicated patterns, again as before. Compared to the septoria model, the boundary between the areas where mixture and alternation perform better for selection curves in the opposite direction; concave upward rather than downward (compare Figure 10a with Figure 4a). This is because the maximum effectiveness of the low-risk in the powdery mildew model is larger than its counterpart in the septoria model (*cf.* analytic predictions from the simple exponential growth model, Methods S4). Again, at full dose of both fungicides the superior strategy for both resistance management and long-term yield is Alternation Low-High.

**Figure 10.**
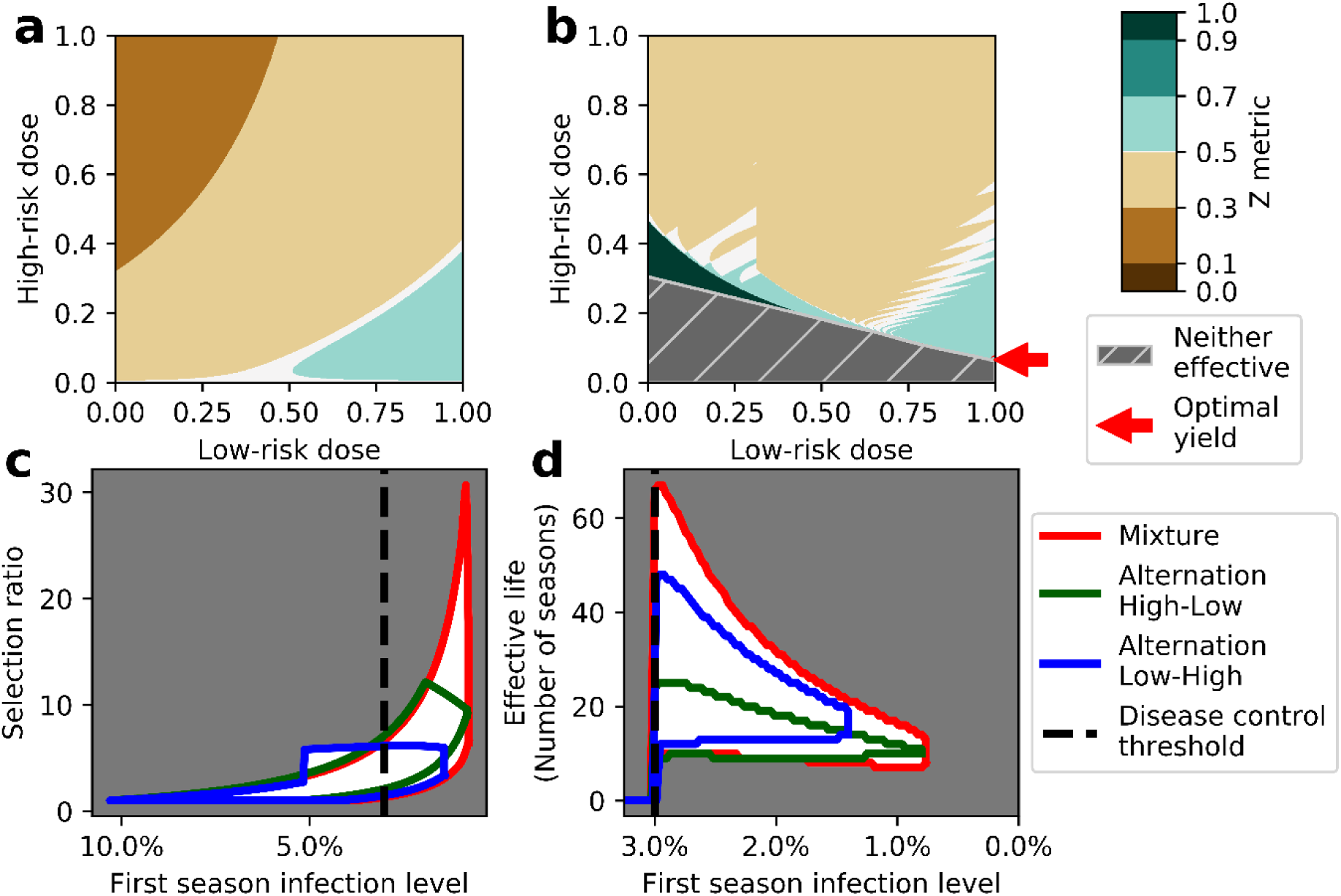
Effect of pathosystem: results for grapevine powdery mildew. (a) Comparison of the selection ratio for mixtures *vs.* the best-performing alternation strategy for the model of powdery mildew on grapevine, for different doses of high-risk and low-risk fungicide (*cf.* Figure 4a, which shows the corresponding result for septoria on winter wheat). (b) Corresponding comparison of lifetime yield in dose-space (*cf.* Figure 4b). (c) Range of selection ratios as a function of the first season infection level for the grapevine model (*cf.* Figure 5f). (d) Range of lifetime yields as a function of the first season infection level for the grapevine model (*cf.* Figure 5g). Again (*cf.* Figure 5f and g), ranges are almost always wider for mixtures than for either alternation strategy, meaning that – depending on the dose of high-risk fungicide – mixtures can lead to both better and worse outcomes for resistance management at any level of disease control. However, dose combinations that cause mixtures to provide the most effective resistance management can always be selected. The dashed lines in (c) and (d) show the minimal acceptable level of disease control.

However, when normalising strategies by the level of initial control (Figures 10c and 10d), mixtures are again capable of generating lower selection pressures and higher effective lives. Compared to septoria, the alternation strategies have less overlap in their performance and the worst-yielding mixture strategies are much more similar to the worst-yielding alternation. Nevertheless, the key result is that the overall optimal strategy for long-term yield is – yet again – to apply as much low-risk as possible, combined with slightly more high-risk than needed for an acceptable initial level of disease control, and to do so under mixture.

## DISCUSSION

We considered resistance management of a fungicide at high-risk of resistance, comparing performance of combining the high-risk chemical with a low-risk fungicide sprayed as either a mixture or in alternation. We assessed performance via the lifetime yield before control breaks down due to fungicide resistance, performing four distinct comparisons. The simplest comparison considered full label doses of both chemicals. For septoria, the largest lifetime yield was obtained by spraying in alternation, although the improvement relative to mixture was relatively small (Figures 3 and 4a and 4b). Alternation was also optimal at full doses in our model of grapevine powdery mildew (Figures 10a and 10b). While performance at full doses of both chemicals is a simple comparison, it is now common practice in some countries for fungicides to be used at lower doses (Jørgensen *et al.*, 2017). Additionally, the comparison depends strongly on model parameterisation. By altering values of epidemiological and/or fungicide-performance parameters, either mixtures or alternation can optimise lifetime yield at full dose (*cf.* changes in colour along the top of individual panels in Figure 7). The set of mechanisms included in the underlying epidemiological model can also affect whether alternation or mixture is the best strategy (note how closely regions shaded light-green approach the points at which *C_H_* = *C_L_* = 1.0 in some panels of Figure 9). Results of the full dose comparison are therefore equivocal, being system-, model-, and parameter-specific. There is also no guarantee that larger lifetime yields would not be obtained by spraying smaller amounts of fungicide, due to the smaller amount of selection that would thus be exerted.

Our second comparison therefore considered performance across all pairs of permissible doses. There are regions of dose-space within which alternations and mixtures each optimise lifetime yield (Figure 4b). The broad pattern in dose-space is robust to model structure (Figures 8 and 9) and pathosystem (Figures 10a and b). An underlying driver of the variation in performance at different doses is – when normalising by the applied doses – that different strategies lead to varying levels of disease control in the absence of resistance (Figures 4c and 5a). An alternate normalisation – as in our third comparison – accounts for this by selecting combinations of pairs of fungicide doses for each strategy that lead to identical initial levels of control (Figure 5f and 5g). The effective lifetime and so lifetime yield for any given level of disease control then depends strongly on the amount of high-risk fungicide that is sprayed (Figure 5h). Since the low-risk chemical exerts no selection in our model, it is optimal to include as much low-risk fungicide as possible in any spray programme, and to combine this with as little high-risk fungicide as provides effective control (Figures 4b and 10b). Arguably this is unsurprising (Shaw, 2006), but focuses our attention on identifying the strategy which has the longest effective lifetime, and so the largest lifetime yield.

For this fourth and final comparison, for both pathosystems we considered (Figures 4b and 10b), and for all model parameterisations (Figures 6 and 7) and sets of epidemiological mechanisms (Figure 9) we tested, the maximum effective lifetime was obtained when fungicides were sprayed as a mixture. We found it was then always optimal to apply a full dose of low-risk mixed with close to the minimal dose of high-risk that retains effective control when there is no resistant pathogen. This optimises the lifetime yield (red arrows in Figures 4b and 10b, and red lines in Figure 7). This spray programme represents the true optimum of all strategies we considered.

Mathematical models of whether alternations or mixtures are better for fungicide-resistance management have been developed for decades. In the early literature whether mixtures provided any benefit beyond allowing a reduction in dose was contentious, and without explicit consideration of dose-response curves, the cost of dose-splitting was not obvious. The recent formalisation of the governing principles has allowed us to clearly disentangle the mechanisms driving the effect of mixtures on selection. The other major distinguishing feature of our work is our extensive sensitivity analysis to model parameterisation as well as the pathosystem that is modelled. We also tested – via our structural sensitivity analysis to model structure – how the set of epidemiological mechanisms included in the underlying model affected our conclusions. It is remarkable that the optimum over all strategies was independent of all of these factors.

However, arguably the most important aspect of our work is that we have provided a concrete explanation for our observations, showing how the governing principles introduced by Milgroom & Fry (1988) and more recently formalised by van den Bosch *et al.* (2014a) can explain the behaviour. The driving mechanism underpinning the relative success of mixtures is that dose-splitting of the low-risk fungicide gives better background disease control and permits a lower dose of high-risk to be used. The growth rate of the pathogen when the high-risk is applied is also suppressed by the low-risk, reducing selection further. For the set of models and parameterisations tested here these effects outweigh the negative effect of the high-risk chemical spraying twice as often. While we have not – and in general cannot – prove this will happen in all parameterisations of all models of all structures for all pathosystems, taken collectively our results provide very good evidence that applying fungicides as mixtures will be the best resistance management strategy in a range of situations. Furthermore, for a set of 1000 parameter sets in which each parameter was sampled uniformly at random from the ranges shown in Figure 7, no case was found where mixture did not provide an overall better lifetime yield than alternation. (Methods S5).

The majority of our results are explained by dose-splitting and suppression by the mixing partner, both of which are simply explained by the governing principles. However, the fact that the two alternation strategies do not perform identically – and that these two strategies differ only in the order in which high-risk and low-risk chemicals are applied – shows that timing of fungicide application can also be important. These can likely also be explained by the governing principles, but with greater difficulty due to the non-trivial interactions between the end of the season, the growth rate of the pathogen at any given time, and the critical period for yield formation. We have therefore not pursued these differences here.

While we have identified how the optimal strategy and combination of doses could be selected, there are potentially issues in adopting our prescribed strategy. The first difficulty is that it requires the threshold between effective and ineffective control to be unambiguously identified. Given the high level of year-on-year variability typical of real disease systems (te Beest *et al.*, 2008) and the extent to which available models do not necessarily capture the complex dynamics of epidemics accurately enough to make such a precise prediction (Gent *et al.*, 2013), this might be rather difficult in practice. There would also be questions raised surrounding the risk-aversion of growers and/or agronomists, who might – reasonably enough – wish to use higher doses of fungicides than are necessary on average to avoid failure of control in years with high disease pressures (Jørgensen *et al.*, 2017), although in principle this might be mitigated via a sufficiently well-calibrated decision support system (Carisse *et al.*, 2010). We have also not considered the economic aspects of our recommendations (te Beest *et al.*, 2013), nor the potentially confounding effects of varying the timing of fungicide sprays (van den Berg *et al.*, 2013, 2016), nor the emergence phase of resistance (Hobbelen *et al.*, 2014; Mikaberidze *et al.*, 2017), nor of spatial heterogeneity in coverage (Shaw, 2000; Parnell *et al.*, 2005, 2006). Nevertheless, by showing in detail and for the first time how fungicide anti-resistance strategies should properly be compared, as well as by showing how results of such comparisons can be explained using simple and intuitive epidemiological principles, our work has developed a firm base to which these complexities can be added. Our future work will do this, albeit with the expectation that mixtures will very often be the better strategy.

## Acknowledgements

JADE acknowledges the BBSRC for support via a University of Cambridge DTP PhD studentship. Rothamsted Research receives support from the Biotechnology and Biological Sciences Research Council (BBSRC) of the United Kingdom.

## Author contributions

JADE, FvdB and NJC conceived and designed the research. JADE performed the experiments and analysed the data. JADE and NJC wrote the paper, with input from FvdB and FJL-R.

## SUPPORTING INFORMATION

### Methods S1 Model of septoria leaf blotch on winter wheat

As described in the main text, the model of septoria leaf blotch on winter wheat is a semi-discrete, compartmental model that runs over successive growing seasons. The model was derived, parameterised and tested against field data in Hobbelen *et al.* (2011).

The model tracks the leaf area index (LAI), the area of leaf per area of ground, of different classes of leaf tissue, distinguishing a number of epidemiologically-relevant compartments: the area of healthy uninfected tissue (**<u>S</u>**usceptible), the area of latent (**<u>E</u>**xposed) and infectious (**<u>I</u>**nfectious) lesions, and the area of dead tissue (**<u>R</u>**emoved). All classes involving the pathogen are divided into separate sub-compartments for the fungicide-resistant (subscript R) and fungicide–sensitive (subscript S) strains. Note that the fungicide-resistant and fungicide-sensitive strain dynamics are identical except that the high-risk fungicide does not affect the fungicide-resistant strain.

We denote the total upper leaf LAI as *A*, with

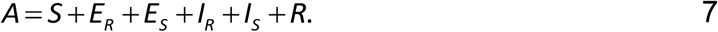

This LAI grows at rate *g*, which is monomolecular after the emergence of the first leaf tracked, and in which disease has no effect on growth

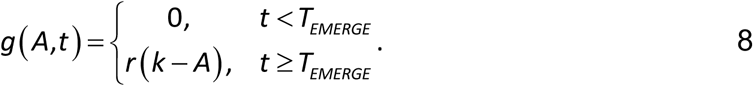

Living host tissues senesce at a rate Γ governed by the time in the season relative to key growth stages,

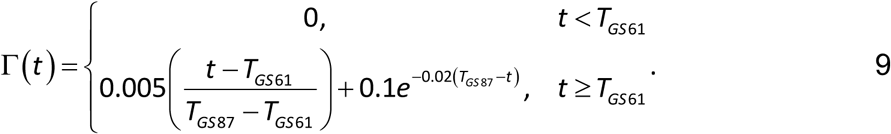

The system of differential equations describing the system is then

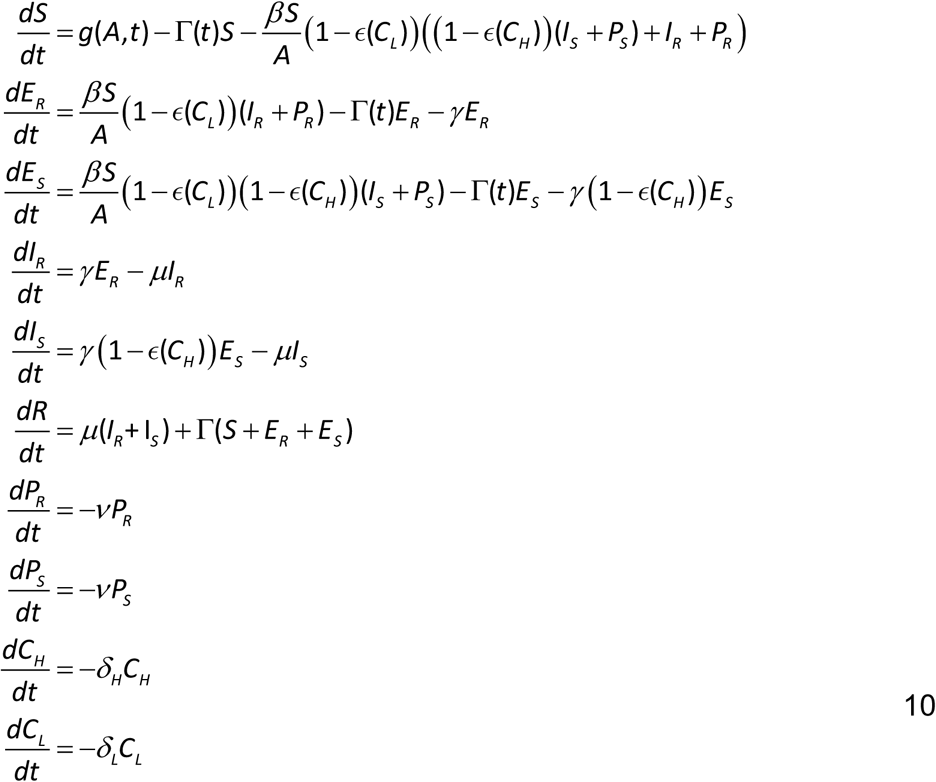

The model was slightly updated relative to the original publication (Hobbelen *et al.* 2011) by modelling disease spread on the top 5 rather than the top 3 leaves. This avoided an edge effect whereby the spray at GS32 (very near to the start of the growing season as modelled in the Hobbelen paper) exerted an unrealistically large amount of selection. This was a modelling artefact due to extremely large per-capita growth rates at the start of the modelled season (caused by primary infection from the time-decaying inoculum), which was fixed by shifting the effective start of the growing season back by two phyllochrons. This change additionally facilitates comparison with later models which track these additional leaves (van den Berg *et al.* 2013). This required the infection rate (β) be refitted, which was done by minimising the squared difference between areas of infectious tissue, summed over every degree-day, between the Hobbelen *et al.* (2013) model and the new parameterisation starting at the emergence of leaf 5. The optimisation was carried out for the times from the emergence of leaf 3 onwards and with no fungicide applied.

### Methods S2 Sensitivity analysis to model structure for the model of septoria leaf blotch

As described in the main text, we investigate the effect of three main features of the septoria model: host-limited infection, whether the latent period is modelled, and phenology. Each of these features corresponds to certain features being included/omitted from the model

- **Host-limited infection**. In models which do not include host-limited infection, the infection rate is independent of the amount of host tissue, and so the terms for infection in Equation 10 are altered as follows

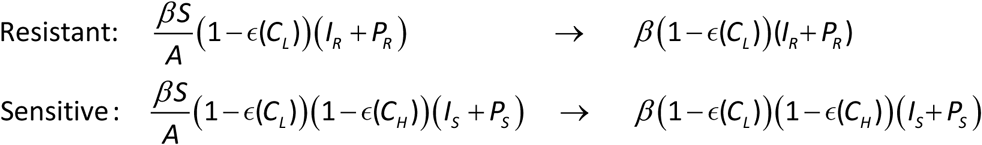
- **Latent period**. If the latent period is removed, infection moves tissue directly from the *S* class to *I*_R_ and *I*_S_ without passing through *E*_R_ and *E*_S_.
- **Phenology**. If phenology is removed from the model, the senescence term Γ(t) (Equation 9) is set to zero and the state variables corresponding to primary inoculum (P_r_ and P_S_) are removed from the model. The epidemic is then started each season by adding a small amount of E_R_ and E_S_ (or I_R_ and I_S_ if there is also no latent period in the simplified model). The amount added is the same as the amount of primary inoculum that would have been present if phenology were included in the model.

Every possible model which either includes or excludes each of these three factors is considered, leading to a total of eight different models. Since models without host-limited infection cannot provide information about the loss of green tissue to infection, they are excluded for the yield analysis, leaving 4 models in that case.

The infection rate parameter (β) is refitted for each model, to allow results to be directly compared. The fitting was done by generating data for every degree-day from the full model when spraying under each of the spraying strategies (mixture and the two alternations) at four different doses of each fungicide (0.25, 0.5, 0.75 and full dose) and when there is no fungicide-resistant pathogen. The value of the infection rate parameter for each simplified model was chosen that minimised the sum of squared differences between the curves for the amount of I_S_ over a single season for the full model and the given simplified model. Multiple doses and strategies were used in the fitting in order to give a parameter value that gave overall similar dynamics across the range of doses that were compared. The results of this fitting are given in Figure S1.

### Methods S3 Model of powdery mildew on grapevine

Similarly to the model of winter wheat septoria leaf blotch, the model of powdery mildew on grapevine is a semi-discrete, compartmental model and runs over multiple growing seasons. The model was derived and parameterised in Burie *et al.* (2011).

The model tracks: healthy uninfected tissue (**<u>S</u>**usceptible), the area of latent (**<u>E</u>**xposed) and infectious (**<u>I</u>**nfectious) lesions, leaf area that has developed resistance to disease due to age (**<u>O</u>**ntogenic) and dead tissue (**<u>R</u>**emoved). All classes involving the pathogen are divided into a sub-compartment for the fungicide-resistant (subscript R) and fungicide-sensitive (subscript S) strains. Again the fungicide-resistant and fungicide–sensitive strain dynamics are almost identical except that the high-risk fungicide does not affect the fungicide-resistant strain.

The total leaf area is

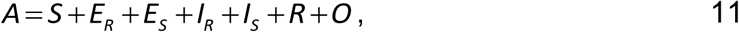

and this is assumed to grow logistically, with

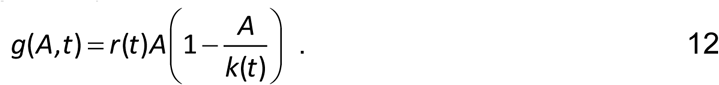

The growth parameters *r*(t) and *k*(t) are piecewise constant functions depending on whether *t* is before or after shoot topping, which is the agronomic practice of removing the upper shoots to encourage secondary growth (see also Table S2). Shoot topping is modelled as occurring on day 173 of the season, and changes the value of these host growth parameters, as well as the infection rate parameter. It also leads to a 20% reduction in the size of the state variable for each compartment.

The system of ODEs describing the powdery mildew model is

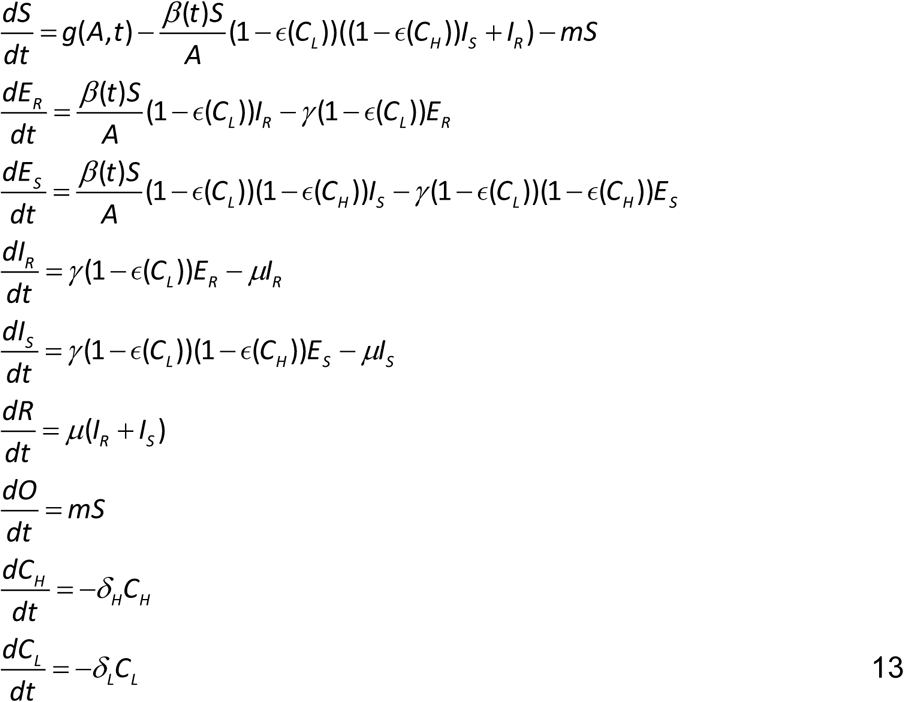

The values for the fungicide dose-response and decay parameters were matched to data from the literature. Jyot *et al.* (2010) measured half-lives of around 3 days for trifloxystrobin on grapevine, and Nasr (2010) found half-lives of around 4 days for sulphur on tomatoes and squashes. Little data was available was available on fungicide effectiveness and so the assumption was made that the maximum effectiveness of both fungicides was 1, representing the assumption that given a suitably high dose of either fungicide the growth of the pathogen can be almost entirely suppressed (even if only for a short time). Reuveni *et al.* (2001) provides the reduction in disease severity when 6 sprays of sulphur or trifloxystrobin were used. The model was set up so that 6 sprays of fungicide were applied starting at day 120, with 14 days between sprays. The values for the curvatures of both fungicides that minimised the sum of squared differences between the percentage reduction in disease severity (compared to untreated) in the model and in the 1999 dataset from Reuveni *et al.* (2001) was then calculated, summing values from each individual day in the models’ results.

### Methods S4 Analysis of selection in a very simple model of resistance dynamics

We consider here an extremely simple model which allows the cumulative selection coefficient – under both mixture and alternation – to be calculated analytically. We ignore exponential decay of fungicides, and instead assume that fungicides remain at fixed dose for a certain amount of time after spraying. There is no unique way to map the applied doses when there is no decay to that case when fungicides decay that conserves the total effect of the fungicides (i.e. integral with respect to time), since the mapping depends on the timescale. Instead the dose applied when there is no decay is simply assumed to be the same as with decay. This final simplification leads to a simple exponential model, of a form very similar to those of the early fungicide resistance modelling literature (Kable and Jeffery 1980; Skylakakis 1981)

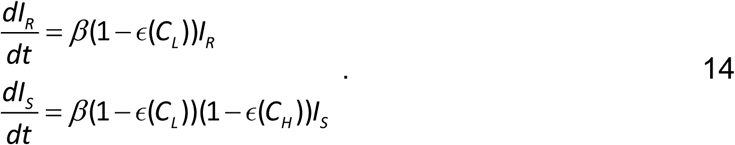

Taking the selection coefficient to be the difference in per capita growth rates of the two strains, and assuming that the fungicides are present for twice as long under mixture as under alternation leads to an analytical form for the cumulative selection coefficients under each strategy:

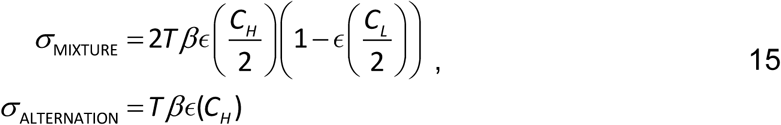

in which *T* is the time of exposure to fungicide under alternation, and where the additional factor of 2 for mixture is because fungicide is then sprayed twice as often.

The ratio of these two quantities quantifies whether mixture (ratio greater than 1) or alternation (ratio less than 1) provides better resistance management

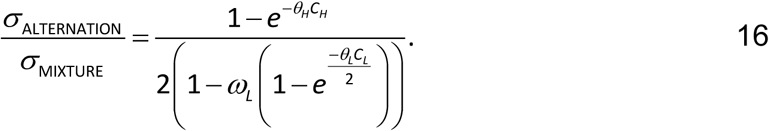

Note that the underlying infection rate (β), the maximum effect of the high-risk fungicide (ω_H_), and the time for which fungicide is present (*T*) all cancel out of this expression.

If we examine the case where this ratio is equal to 1, and ignore the solution *C*_H_ = 0, we can calculate the equation of the boundary curve between the area where mixture and alternation perform better

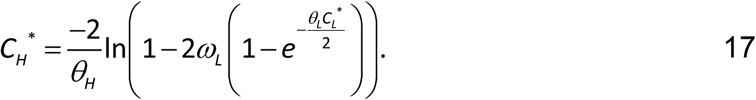

Taking the derivative with respect to the low-risk fungicide dose at equal performance

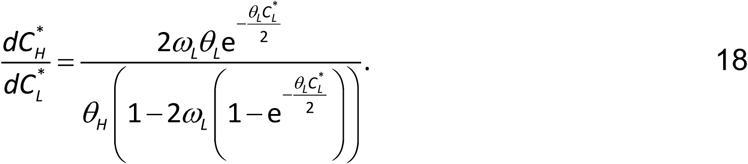

Given that all parameters are strictly positive then the sign of this equation is determined by the sign of the denominator which is positive if and only if ε(*C*_L_^*^/2) > ½. However, if we look at the equation for the boundary curve, *C*_H_ ^*^ is undefined if this condition is met, since Equation 17 is infinite for ε(*C*_L_ ^*^/2) = ½. As the first derivative cannot be zero there are no turning points and the boundary function is monotonic. We can conclude that are always areas where both strategies can out-perform the other (although they can be small) separated by a monotonically-increasing boundary of equal performance.

We can also examine the sign of the second derivative of the boundary function with respect to the equilibrium low-risk fungicide dose. If it is always positive then the curve is convex in (*C*_L_ ^*^,*C*_H_ ^*^) space; if negative, then the curve is concave; and if zero, then the curve is simply a straight line. Since

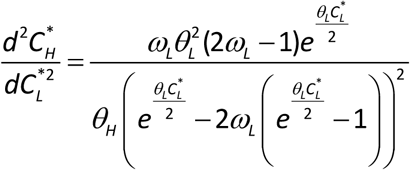

The sign of the equation is controlled by ω_L_ alone. If less than ½ the boundary is concave, greater convex and if equal straight. This is because the value of ω_L_ controls whether the low-risk can make up for the increased effect of the high-risk under mixture, in the limit of large amounts of both fungicides. Considering the value of Equation 16 for *C*_H_=1 and *C*_L_=0 we see that alternation is always superior at this point, therefore alternation will always perform better above the boundary curve and mixtures below.

### Methods S5 Testing robustness of the result that mixture outperforms alternation

In order to provide further evidence that the superior performance of mixtures was not specific to our chosen parameterisation of the septoria model we carried out the following additional test.

- Repeat until 1000 parameter value sets are accepted
  - Choose parameter values uniformly at random from within the ranges of all parameters displayed in Figure 7 (note this means that all parameters will in general take non-default values)
  - Check if parameters give realistic solutions and continue if so, otherwise generate new parameters. For mixtures and both alternation strategies the reality check consists of ensuring:
    - that yield is below 95% when the dose of high-risk is zero and the dose of low-risk is one;
    - that yield is above 95% when a full dose of both high-risk and low-risk are applied.
  - At full dose of the low-risk, find the optimal dose of high-risk and the application strategy that gives the largest lifetime yield

As stated in the main text, no case was found which led to alternation out-performing mixture. The volume of parameter space giving realistic solutions was high, in general only requiring one or two attempts at parameter value generation to identify a reasonable set of parameters (i.e. approximately 50% of parameters tested lead to a realistic parameterisation of the model).

**Figure S1.**
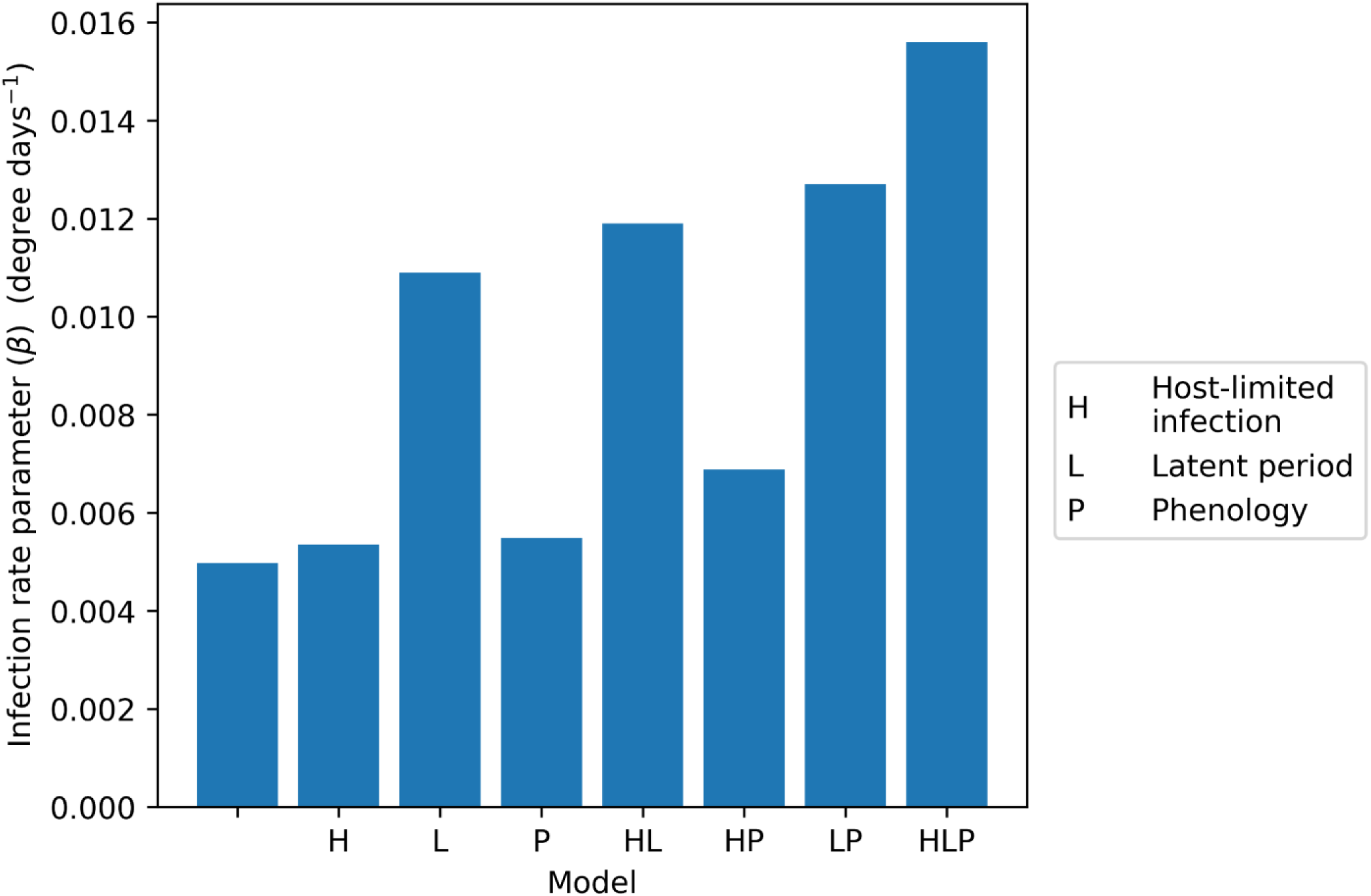
Fitted values of the infection rate parameters in the simpler models including fewer epidemiological mechanisms which are used in the sensitivity analysis to model structure. The set of mechanisms included in each model is specified by the combination of letters H, L and P corresponding to host-limited infection, latent period and phenology, respectively (see main text). The HLP model is therefore the full model considered in the majority of the main text; the other models appear in Figures 8 and 9. The values of infection rate parameters were found by fitting the simpler models to the results of the more complex full model at a range of fungicide doses (the procedure for doing this is fully described in Methods S2).

**Figure S2.**
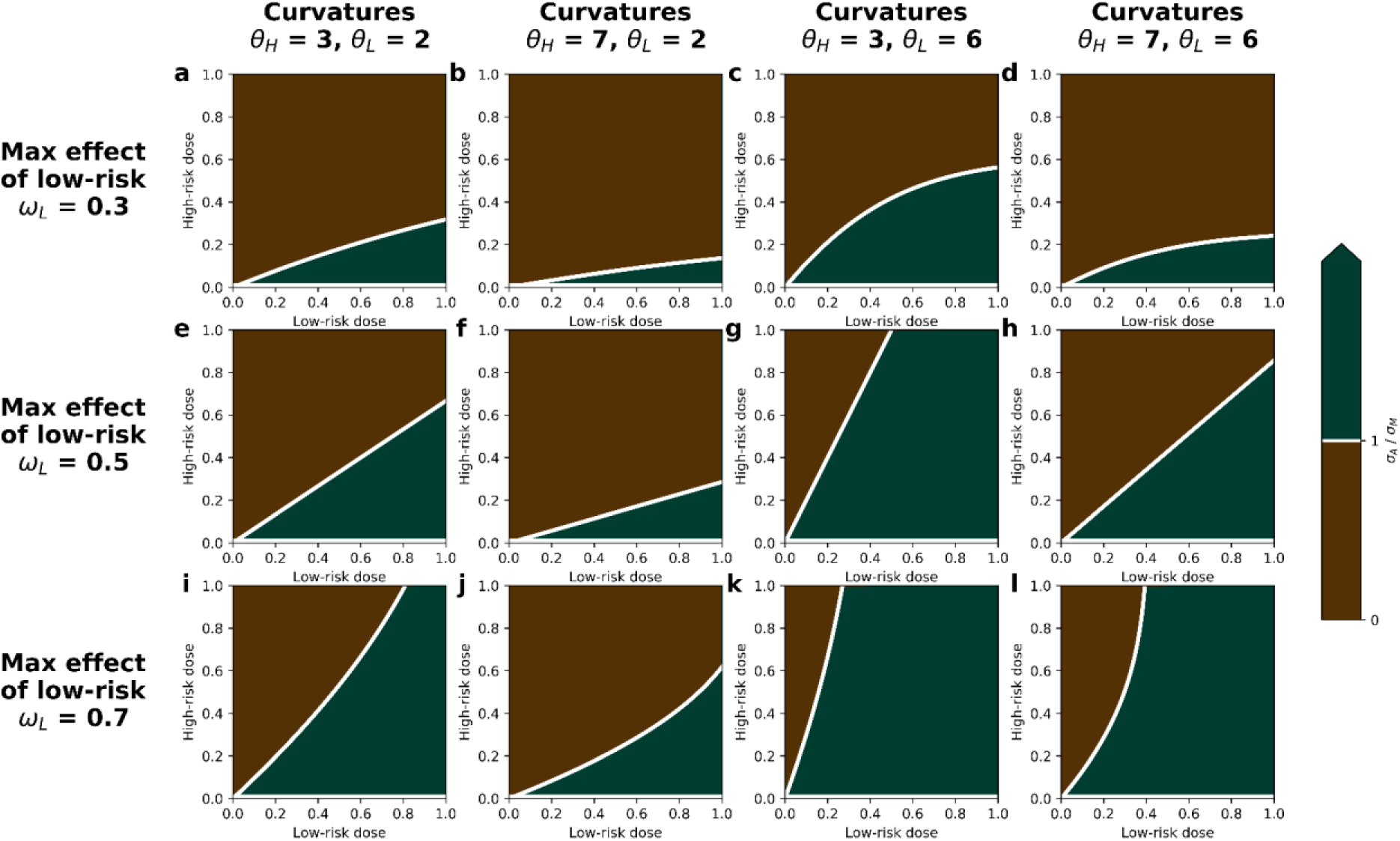
Selection in dose space for a simple exponential growth model with no decay of fungicides, showing whether mixtures or alternation have a smaller cumulative selection coefficient (σ, the time integral of the selection coefficient). Green areas show regions within which mixture performs better, and brown shows regions within which alternation is better. Responses are shown for the simple exponential growth model with non-decaying fungicides (Equation 14; Methods S4). The analytic prediction indicates that in this model the maximum effect of the high-risk fungicide (ω_H_) has no effect on relative strategy performance; neither does the underlying pathogen growth rate (β). The values of the other fungicide parameters – i.e. the maximum effect of low-risk fungicide (ω_L_), and the curvature parameters of both fungicides (θ_H_ and θ_L_) – therefore control the shape of the response. The first row (**a-d**) has ω_L_ = 0.3; the second row (**e-h**) has ω_L_ = 0.5; and the third row (**i-l**) has ω_L_ = 0.7. The first column (**a, e, i**) has θ_H_ = 3 and θ_L_ = 2, the second (**b, f, j**) has θ_H_ = 7 and θ_L_ = 2, the third (**c, g, k**) has θ_H_ = 3 and θ_L_ = 6, and the fourth (**d, h, l**) has θ_H_ = 7 and θ_L_ = 6.

**Table S1.** Default parameterisation of the model of septoria leaf blotch on winter wheat.

| Parameter | Symbol | Value | Reference |
| --- | --- | --- | --- |
| Infection rate parameter | $\beta$ | $1.56 \times 10^{-2}$ degree-day<sup>-1</sup> | Fit to output of model of Hobbelen *et al.* 2013 (see Methods S1) |
| Latent period | $1/\gamma$ | 266 degree-days | Hobbelen *et al.* 2013 |
| Infectious period | $1/\mu$ | 456 degree-days | Hobbelen *et al.* 2013 |
| High risk fungicide name |  | Pyraclostrobin | Hobbelen *et al.* 2011 |
| High risk fungicide activity |  | Protectant / Eradicant | Hobbelen *et al.* 2011 |
| High risk fungicide maximum effect | $\omega_H$ | 1 | Hobbelen *et al.* 2011 |
| High risk fungicide dose response curvature | $\theta_H$ | 9.6 | Hobbelen *et al.* 2011 |
| High risk fungicide decay parameter | $\delta_H$ | $1.11 \times 10^{-2}$ degree-days<sup>-1</sup> | Hobbelen *et al.* 2011 |
| Low risk fungicide name |  | Chlorothalonil | Hobbelen *et al.* 2011 |
| Low risk fungicide activity |  | Protectant | Hobbelen *et al.* 2011 |
| Low risk fungicide maximum effect | $\omega_L$ | 0.48 | Hobbelen *et al.* 2011 |
| Low risk fungicide dose response curvature | $\theta_L$ | 9.9 | Hobbelen *et al.* 2011 |
| Low risk fungicide decay parameter | $\delta_L$ | $6.91 \times 10^{-3}$ degree-days<sup>-1</sup> | Hobbelen *et al.* 2011 |
| Initial resistance frequency | $\psi$ | $1 \times 10^{-10}$ | Assumed (a range of values were tested; see Figure 7h in main text) |
| Initial inoculum density | $\phi$ | $1.09 \times 10^{-2}$ | Hobbelen *et al.* 2011 |
| Initial area of susceptible leaf | $S(0)$ | 0.05 | Hobbelen *et al.* 2011 |
| Initial area of leaf with latent infection by susceptible strain | $E_S(0)$ | 0 | Hobbelen *et al.* 2011 |
| Initial area of leaf with latent infection by resistant strain | $E_R(0)$ | 0 | Hobbelen *et al.* 2011 |
| Initial area of leaf infected by susceptible strain | $I_S(0)$ | 0 | Hobbelen *et al.* 2011 |
| Initial area of leaf infected by resistant strain | $I_R(0)$ | 0 | Hobbelen *et al.* 2011 |
| Initial area of dead leaf | $R(0)$ | 0 | Hobbelen *et al.* 2011 |
| Host growth rate parameter | $r$ | $1.26 \times 10^{-2}$ degree-days<sup>-1</sup> | Hobbelen *et al.* 2011 |
| Host carrying capacity | $k$ | 4.2 | van den Berg *et al.* 2013 |
| Decay rate of primary inoculum | $\nu$ | $8.5 \times 10^{-3} \text{ degree-days}^{-1}$ | Hobbelen *et al.* 2011 |
| Spray times |  | GS32, GS39 | Hobbelen *et al.* 2014; Paveley *et al.* 2014 |
| Time of emergence of leaf 5 | $T_{\text{EMERGE}}$ | 1212 degree days | van den Berg *et al.* 2013 |
| Time of GS32 | $T_{\text{GS32}}$ | 1456 degree days | van den Berg *et al.* 2013 |
| Time of GS39 | $T_{\text{GS39}}$ | 1700 degree days | van den Berg *et al.* 2013 |
| Time of GS61 | $T_{\text{GS61}}$ | 2066 degree days | van den Berg *et al.* 2013 |
| Time of GS87 | $T_{\text{GS87}}$ | 2900 degree days | van den Berg *et al.* 2013 |
| Initial area of lower leaf infected by resistant strain | $P_R(0)$ | $\psi\phi$ | Derived |
| Initial area of lower leaf infected by susceptible strain | $P_S(0)$ | $(1-\psi)\phi$ | Derived |

**Table S2.** Default parameterisation of the model of powdery mildew on grapevine

| Parameter | Symbol | Value |  | Reference |
| --- | --- | --- | --- | --- |
| Infection rate parameter | $\beta$ | $1.605 \text{ cm}^{-2} \text{ day}^{-1}$ | $1.688 \text{ cm}^{-2} \text{ day}^{-1}$ | Burie *et al.* 2011 |
| Latent period | $1/\gamma$ | 10 days | | Burie *et al.* 2011 |
| Infectious period | $1/\mu$ | 10 days | | Burie *et al.* 2011 |
| High risk fungicide name |  | Trifloxystrobin |  |  |
| High risk fungicide activity |  | Protectant / Eradicant |  |  |
| High risk fungicide maximum effect | $\omega_H$ | 1 | | Assumed |
| High risk fungicide dose response curvature | $\theta_H$ | 4.88 | | Fitted to data from Reuveni 2001 (see Methods S3) |
| High risk fungicide decay parameter | $\delta_H$ | 0.231 | | Taken from range of values in Jyot *et al.* 2010 (see Methods S3) |
| Low risk fungicide name |  | Sulphur |  |  |
| Low risk fungicide activity |  | Protectant / Eradicant |  |  |
| Low risk fungicide maximum effect | $\omega_L$ | 1 | | Assumed |
| Low risk fungicide dose response curvature | $\theta_L$ | 1.02 | | Fitted to data from Reuveni 2001 (see Methods S3) |
| Low risk fungicide decay parameter | $\delta_L$ | $0.173 \text{ days}^{-1}$ | | Taken from range of values in Nasr 2010 (see Methods S3) |
| Initial resistance frequency | $\psi$ | $1 \times 10^{-10}$ | | Assumed |
| Initial inoculum density | $\phi$ | $0.13 \text{ cm}^2$ | | Burie *et al.* 2011 |
| Initial area of susceptible leaf | $S(0)$ | $42.34 \text{ cm}^2$ | | Burie *et al.* 2011 |
| Initial area of leaf with latent infection by susceptible strain | $E_S(0)$ | $(1-\psi) \phi$ | | Derived |
| Initial area of leaf with latent infection by resistant strain | $E_R(0)$ | $\psi \phi$ | | Derived |
| Initial area of leaf infected by susceptible strain | $I_S(0)$ | $0 \text{ cm}^2$ | | Burie *et al.* 2011 |
| Initial area of leaf infected by resistant strain | $I_R(0)$ | $0 \text{ cm}^2$ | | Burie *et al.* 2011 |
| Initial area of dead leaf | $R(0)$ | $0 \text{ cm}^2$ | | Burie *et al.* 2011 |
| Host growth rate parameter | $r$ | $0.147 \text{ days}^{-1}$ | $0.032 \text{ days}^{-1}$ | Burie *et al.* 2011 |
| Host carrying capacity | $k$ | 26106<br>cm<sup>2</sup> | 2.461 x<br>10<sup>8</sup> cm<sup>2</sup> | Burie *et al.* 2011 |
| Spray times |  | Days 161, 174 |  | Assumed |
| Initial area of<br>ontogenically resistant<br>leaf | $O(0)$ | 0 cm<sup>2</sup> | | Burie *et al.* 2011 |
| Rate of ontogenic<br>resistance development | $m$ | 0.1 days<sup>-1</sup> | | Burie *et al.* 2011 |
| Time of end of<br>simulation | $T_{END}$ | Day 230 | | Assumed |
| Time of primary<br>infection | $T_{INF}$ | Day 119 | | Mammeri *et al.*, 2014 |
| Time of flowering | $T_{FLO}$ | Day 163 | | Mammeri *et al.*, 2014 |
| Time of shoot topping | $T_{TOP}$ | Day 173 | | Mammeri *et al.*, 2014 |
| % leaf area lost on<br>topping |  | 20% |  | Burie *et al.* 2011 |

